# Cell-specific Bioorthogonal Tagging of Glycoproteins

**DOI:** 10.1101/2021.07.28.454135

**Authors:** Anna Cioce, Beatriz Calle, Tatiana Rizou, Sarah C. Lowery, Victoria Bridgeman, Keira E. Mahoney, Andrea Marchesi, Ganka Bineva-Todd, Helen Flynn, Zhen Li, Omur Y. Tastan, Chloe Roustan, Pablo Soro-Barrio, Thomas M. Wood, Tessa Keenan, Peter Both, Kun Huang, Fabio Parmeggiani, Ambrosius P. Snijders, Mark Skehel, Svend Kjaer, Martin A. Fascione, Carolyn R. Bertozzi, Sabine Flitsch, Stacy A. Malaker, Ilaria Malanchi, Benjamin Schumann

## Abstract

Altered glycosylation is an undisputed corollary of cancer development. Understanding these alterations is paramount but hampered by limitations underlying cellular model systems. For instance, the intricate interactions between tumour and host cannot be adequately recapitulated in monoculture of tumour-derived cell lines. More complex co-culture models usually rely on sorting procedures for proteome analyses and rarely capture the details of protein glycosylation. Here, we report a strategy termed Bio-Orthogonal Cell line-specific Tagging of Glycoproteins (BOCTAG). Cells are equipped by transfection with an artificial biosynthetic pathway that transforms bioorthogonally tagged sugars into the corresponding nucleotide-sugars. Only transfected cells incorporate bioorthogonal tags into glycoproteins in the presence of non-transfected cells. We employ BOCTAG as an imaging technique and to annotate cell-specific glycosylation sites in mass spectrometry-glycoproteomics. We demonstrate application in co-culture and mouse models, allowing for profiling of the glycoproteome as an important modulator of cellular function.

## INTRODUCTION

Cancer is a multifactorial disease consisting of an interplay between different host and tumour cells. Emulating the complexity of a tumour using cell monoculture is thus incomplete by design, requiring more elaborated co-culture systems or *in vivo* models.^1–3^ Recent years have seen a stark increase in methods to probe the transcriptomes of tumour and host cell populations, respectively, providing some insight into their state within a multicellular conglomerate.^4^ However, the relationship between transcriptome and proteome is still elusive.^5^ In addition, posttranslational modifications (PTMs) heavily impact the plasticity of the proteome. Glycosylation is the most complex and most abundant PTM, but challenging to probe due to the non-templated nature of glycan biosynthesis.^6^ Glycans are generated by the combinatorial interplay of >250 glycosyltransferases (GTs) and glycosidases, mostly in the secretory pathway.^7^ Certain glycoproteins aberrantly expressed in cancer, such as mucins, are approved as diagnostic markers, but their discovery is a particular challenge.^8,9^ This is especially true when *in vivo* or *in vitro* model systems comprise cell populations from the same organism that do not allow distinction of proteomes by amino acid sequence.^10,11^ Methods to study the glycoproteome of a cell type in co-culture or *in vivo* are therefore an unmet need.

Metabolic oligosaccharide engineering (MOE) produces chemical reporters of glycan subtypes.^12^ MOE reagents are membrane permeable monosaccharide precursors modified with chemical tags amenable to bioorthogonal chemistry.^13^ Following incorporation into the glycoproteome, chemical tags are reacted with traceable enrichment handles or fluorophores, for instance by Cu(I)-catalysed azide-alkyne cycloaddition (CuAAC).^14,15^ Many MOE reagents are based on analogues of sugars such as *N*-acetylgalactosamine (GalNAc) that are straightforward to chemically tag by replacing the acetamide with bioorthogonal *N*-acylamides (Fig. 1a). Unmodified GalNAc is normally activated by the biosynthetic GalNAc salvage pathway to the nucleotide-sugar UDP-GalNAc that can follow two major distinct metabolic fates (Fig. 1a).^14,16–18^ First, the 20 members of the GalNAc transferase family (GalNAc-T1…T20) use UDP-GalNAc to form the linkage GalNAcα-Ser/Thr and thereby prime cancer-relevant O-GalNAc glycans.^14,19,20^ Second, epimerisation at the GalNAc C4 position by the UDP-galactose-4-epimerase (GALE) yields UDP-*N*-acetylglucosamine (UDP-GlcNAc) that can be incorporated into different glycan subtypes, for instance Asn-linked N-glycans.^17,18,21^ Certain chemical modifications at the *N*-acyl moiety can render GalNAc analogues recalcitrant to these metabolic processes. For instance, analogues of UDP-GalNAc with long alkyne-containing *N*-acyl substituents are not biosynthesised by the GalNAc salvage pathway and not used as substrates by wild type (WT)-GalNAc-Ts.^18,22–24^ While being a substantial impediment to generating MOE reporters, we realised that overcoming these metabolic roadblocks might enable programmable bioorthogonal glycoprotein tagging. Such a strategy would allow for studying the glycoproteome in a cell-specific fashion, which is currently elusive despite the rapid advances in the development of new MOE reagents.

**Fig. 1:**
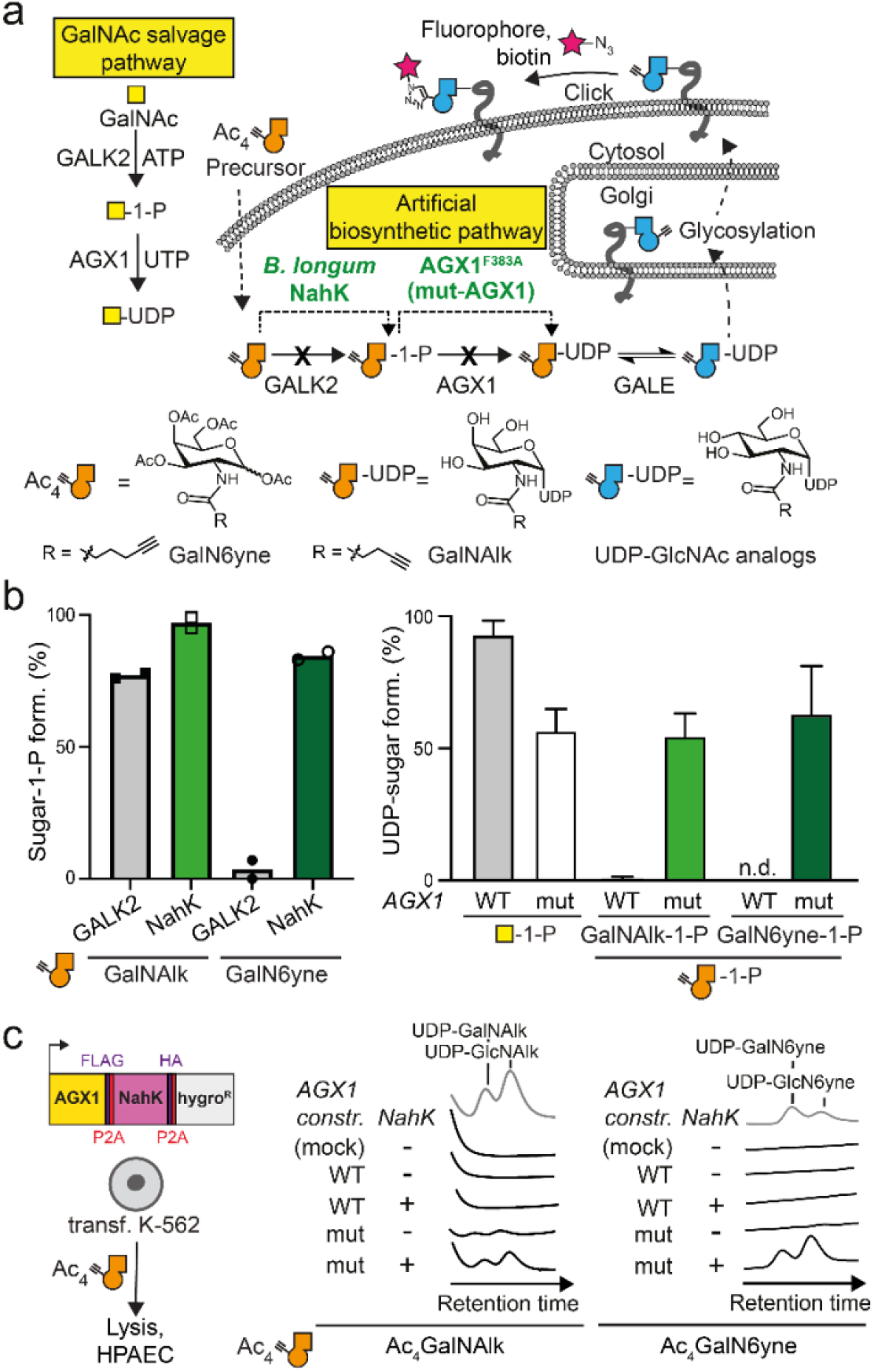
Development of an artificial biosynthetic pathway for chemically tagged UDP-GalNAc/GlcNAc analogues. **a,** strategy of metabolic oligosaccharide engineering. A chemically modified GalNAc analogue that is not accepted by the GalNAc salvage pathway should be processed by an artificial biosynthetic pathway. *B. longum* NahK and mut-AGX1 biosynthesize UDP-GalNAc analogues and, by epimerisation, UDP-GlcNAc analogues. Incorporation into glycoconjugates can be traced by CuAAC. **b,***in vitro* evaluation of GalNAc-1-phosphate analogue synthesis by human GALK2 or *B. longum* NahK (left) and UDP-GalNAc analogue synthesis by WT- or mut-AGX1 (right). Data were recorded in LC-MS assays and processed by integrated ion counts. Data are from two independent experiments and depicted as individual data points and means (left) or from three independent experiments and depicted as means + standard deviation (SD, right). **c,** biosynthesis of UDP-GalNAc/GlcNAc analogues in cells stably expressing both NahK and mut-AGX1 or either component, as assessed by high performance anion exchange chromatography (HPAEC). A hygromycin resistance gene allows for stable transfection. Data are representative of one out of two independent experiments collected on two different days. mock: pSBbi-GH empty plasmid.

Here, we develop a technique called Bio-Orthogonal Cell-specific Tagging of Glycoproteins (BOCTAG). The strategy uses an artificial biosynthetic pathway to generate alkyne-tagged UDP-GalNAc and UDP-GlcNAc analogues from a readily available GalNAc precursor that is not accepted by the GalNAc salvage pathway. We find that a single methylene group between 5-carbon (GalNAlk) and 6-carbon (GalN6yne) *N*-acyl substituents drastically reduces uptake by the native GalNAc salvage pathway and thereby reduces the background of bioorthogonal labelling in non-transfected cells. Only cells carrying the artificial pathway biosynthesise the corresponding UDP-sugars (UDP-GalN6yne and UDP-GlcN6yne) that are then used by GTs to chemically tag the glycoproteome. We further expand the strategy with mutant GalNAc-Ts that are engineered to accept UDP-GalN6yne as a substrate. The combined use of an artificial biosynthetic pathway and engineered GalNAc-Ts enables GalN6yne-mediated fluorescent labelling of the cellular glycoproteome that is two orders of magnitude higher than in cells carrying neither component. We demonstrate that BOCTAG allows for programmable glycoprotein tagging in co-culture and mouse models. Moreover, the nature of the artificial biosynthetic pathway allowed for the use of readily available Ac_4_GalN6yne as a precursor with enhanced stability over previously used caged GalN6yne-1-phosphates as an essential pre-requisite for *in vivo* applications. We show that the chemical modification enters a range of glycan subtypes, supporting the use of BOCTAG to tag a large number of glycoproteins in complex biological systems.

## RESULTS

### Developing an artificial biosynthetic pathway for chemically tagged UDP-sugars

The human GalNAc salvage pathway consists of the kinase GALK2 and the pyrophosphorylases AGX1/2 to convert GalNAc first into GalNAc-1-phosphate and subsequently into UDP-GalNAc, respectively (Fig. 1a). Since neither analogues of GalNAc nor GalNAc-1-phosphate can be utilized by any other metabolic enzyme, the GalNAc salvage pathway was deemed suitable for monitoring conversions of each step while supplying readily accessible synthetic, bioorthogonal precursors. GALK2 and AGX1/2 are impervious to large chemical modifications at the *N*-acyl moiety of GalNAc (Fig. 1a), corroborated by crystal structures of these enzymes (Fig. S1).^18,25–27^ An artificial biosynthetic pathway was thus designed to convert chemically tagged GalNAc analogues first to the corresponding sugar-1-phosphates and subsequently to the UDP-sugars. We chose both a 6-carbon hex-5-ynoate chain (GalN6yne) and a 5-carbon pent-4-ynoate chain (GalNAlk) as GalNAc modifications due to their availability and previous use by us and others.^18,27,28^ In *in vitro* enzymatic assays detected by liquid chromatography-mass spectrometry (LC-MS), recombinant GALK2 accepted GalNAlk as a substrate, but only marginally accepted GalN6yne (Fig. 1b). In contrast, promiscuous bacterial *N*-acetylhexosaminyl kinases (NahK) from various source organisms converted GalN6yne to GalN6yne-1-phosphate almost quantitatively (Fig. 1b, fig. S2a).^29^ Similarly, the pyrophosphorylase AGX1 showed little to no turnover of both GalNAlk-1-phosphate and GalN6yne-1-phosphate to the corresponding UDP-sugars (Fig. 1b). We and others have mutated AGX1 at residue Phe383 to smaller amino acids to accommodate chemical *N*-acyl modifications.^27,30^ AGX1^F383A^, herein called mut-AGX1, converted both synthetic GalNAlk-1-phosphate and GalN6yne-1-phosphate to UDP-GalNAlk and UDP-GalN6yne, respectively (Fig. 1b).

We next assessed UDP-sugar biosynthesis in the living cell. Stable bicistronic expression of a codon-optimized version of *Bifidobacterium longum* NahK as well as mut-AGX1 in K-562 cells biosynthesized UDP-GalNAlk and UDP-GalN6yne from membrane-permeable per-acetylated precursors Ac_4_GalNAlk and Ac_4_GalN6yne, respectively (Fig. 1c). Expression of either enzyme alone or WT-AGX1 led to inefficient biosynthesis compared to levels of native UDP-sugars (Fig. S3). We confirmed these results by feeding cells a caged precursor of GalN6yne-1-phosphate that was uncaged in the living cell and converted to UDP-GalN6yne only in the presence of mut-AGX1 (Fig. S3). Alkyne-tagged UDP-GalNAc analogues were converted to the corresponding UDP-GlcNAc analogues (UDP-GlcNAlk or UDP-GlcN6yne, respectively) in cells by the epimerase GALE, which was corroborated in an *in vitro* epimerisation assay (Fig. 1a,c fig. S2b, fig. S3). Thus, installing an artificial biosynthetic pathway led to programmable biosynthesis of alkyne-tagged analogues of UDP-GalNAc and UDP-GlcNAc.

We next assessed chemical tagging of the cell surface glycoproteome in living cells. K-562 cells stably expressing combinations of NahK and AGX1 were fed with DMSO, Ac_4_GalNAlk or Ac_4_GalN6yne and reacted with the clickable fluorophore CF680-picolyl azide by CuAAC. The MOE reagent Ac_4_ManAlk that enters the pool of the sugar sialic acid was included as a positive control. Alkyne tags were visualized by in-gel fluorescence after cell lysis (Fig. 2a). While Ac_4_GalNAlk feeding led to high-intensity fluorescent signal when NahK and mut-AGX1 were expressed, substantial signal was observed in cells expressing WT-AGX1 when NahK was present (Fig. 2a). Fluorescent signal after Ac_4_GalNAlk feeding was also observed in cells transfected with an empty plasmid or only overexpressing WT-AGX1, confirming the permissiveness of the GalNAc salvage pathway for GalNAlk (Fig. 1b).^18^ In contrast, Ac_4_GalN6yne incorporation was critically dependent on the expression of mut-AGX1, while the presence of NahK led to a further sixfold increase in fluorescence intensity (Fig. 2a). Ac_4_ManAlk gave fluorescent signal regardless of the enzyme combination expressed. Dose response experiments showed that Ac_4_GalN6yne-mediated fluorescence intensity increased over two orders of magnitude with the concentration of the probe between 16 nM and 50 μM only when NahK and mut-AGX1 were present (Fig. 2b). Transfection and feeding with chemically modified sugars can in theory alter the cellular transcriptome, leading to artifacts in protein expression and metabolic labelling. We performed transcriptomic analyses in cells transfected with either NahK/mut-AGX1 or empty plasmid, and fed with either DMSO vehicle, Ac_4_GalN6yne or Ac_4_GalNAc. By performing correlation plot and principal component analysis (PCA)(Fig. S4), we observed that the day of sample collection has a greater effect on transcript levels than either transgene expression or compound treatment (Fig. S4b). These data suggest that neither artificial biosynthetic pathway nor compound feeding has substantial effects on the transcriptome. Due to the robustness of metabolic incorporation, we used Ac_4_GalN6yne as an MOE reagent for all subsequent applications of BOCTAG.

**Fig. 2:**
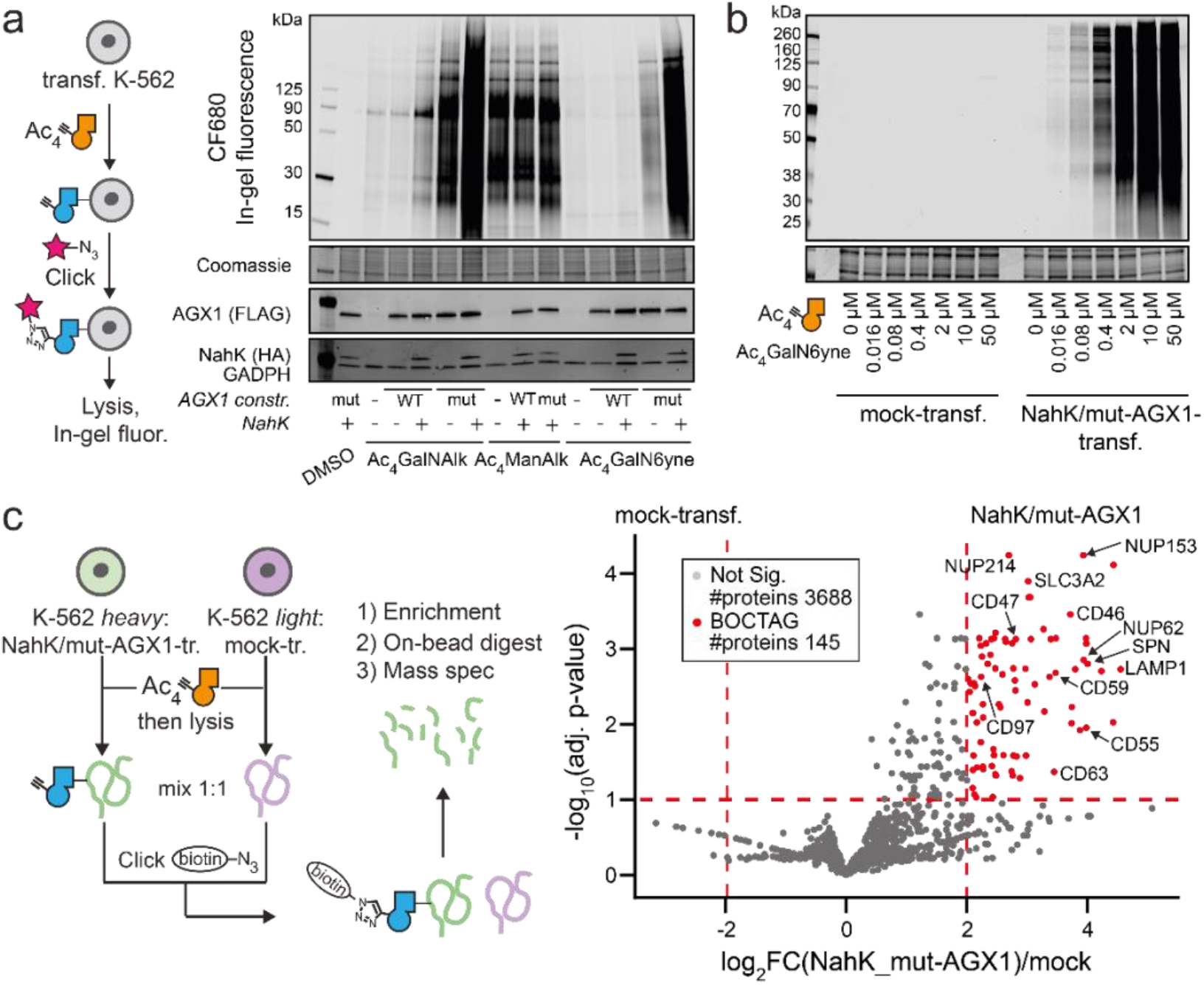
An artificial biosynthetic pathway enables programmable chemical tagging of the glycoproteome. **a,** evaluation of cell surface glycoproteome tagging after treating K-562 cells stably expressing NahK/AGX1 combinations with 50 μM Ac_4_GalNAlk, 50 μM Ac_4_GalN6yne or 10 μM Ac_4_ManNAlk. Glycoproteins were visualised by in-gel fluorescence after treating cells with CF680-picolyl azide under CuAAC conditions and subsequent cell lysis. **b,** dose-response experiment of cell surface glycoproteome tagging, with samples processed as in **a**. Data in **a** and **b** are representative of one out of two independent experiments. **c,** quantitative measurement of glycoprotein tagging by SILAC. Data were analysed from three independent experiments, collected on three different days, with forward (heavy mock, light NahK/mut-AGX1) and reverse (light mock, heavy NahK/mut-AGX1) analyses incorporated as a total of six replicates. Data are visualised as volcano plot, choosing 4-fold enrichment and a p-value of 0.1 as cut-offs, with example glycoproteins annotated. Significance levels were indicated. mock: pSBbi-GH empty plasmid.

### An artificial biosynthetic pathway allows for programmable enrichment of the glycoproteome

We used Stable Isotope Labelling by Amino Acids in Cell Culture (SILAC)-based proteome analysis to confirm and quantify chemical glycoproteome tagging. K-562 cells transfected with NahK/mut-AGX1 or an empty plasmid (mock-transfected). Both were individually grown in heavy or light media in the presence of either Ac_4_GalN6yne or DMSO. Lysates of these cells were mixed as different combinations to contain equal amounts of heavy and light protein, and clickable biotin-picolyl azide was installed on tagged glycoproteins by CuAAC. Enrichment on neutravidin beads followed by on-bead digest allowed analysis by quantitative mass spectrometry (MS). In three independent experiments in which both combinations of heavy and light SILAC labelling each were used (Fig. 2c), we found peptides from 145 proteins to be significantly enriched in NahK/mut-AGX1-transfected cells (Supplementary Table1). More than 99% (143/145) of these proteins have been previously annotated^31–33^ as either N- or O-glycosylated, including the nucleoporins Nup62 and Nup153 and the cell surface proteins CD47 and NOTCH1, confirming the stringency of the approach for tagging glycoproteins.

### Cell type-specific glycoproteome tagging in co-culture

We next assessed the suitability of the artificial biosynthetic pathway NahK/mut-AGX1 as a BOCTAG cell type-specific glycoproteome labelling technique by fluorescence microscopy. Colonies of NahK/mut-AGX1-transfected and GFP-expressing 4T1 breast cancer cells were established on a monolayer of non-transfected MLg fibroblast cells by co-culturing for 72 h before media supplementation with either Ac_4_GalN6yne, Ac_4_ManAlk or DMSO (Fig. 3a). Clickable biotin-picolyl azide was installed by CuAAC followed by Streptavidin-AF647 staining to visualize chemical tagging, and cells were counter-stained with fluorescently labelled phalloidin. Streptavidin-AF647 signal was strongly and reproducibly restricted to GFP-expressing cells only when Ac_4_GalN6yne was fed and NahK/mut-AGX1 were expressed (Fig. 3b, C, fig. S5-S8), indicating a localised BOCTAG signal. In contrast, the promiscuous MOE reagent Ac_4_ManNAlk was non-specifically incorporated throughout the entire co-culture (Fig. 3b, c, fig. S5-S8). When both GFP-4T1 and MLg cell lines expressed NahK/mut-AGX1, both exhibited a strong Streptavidin-AF647 signal (Fig. S8b). Taken together, BOCTAG enables cell-specific tagging of cell surface glycoproteins in co-culture.

**Fig. 3:**
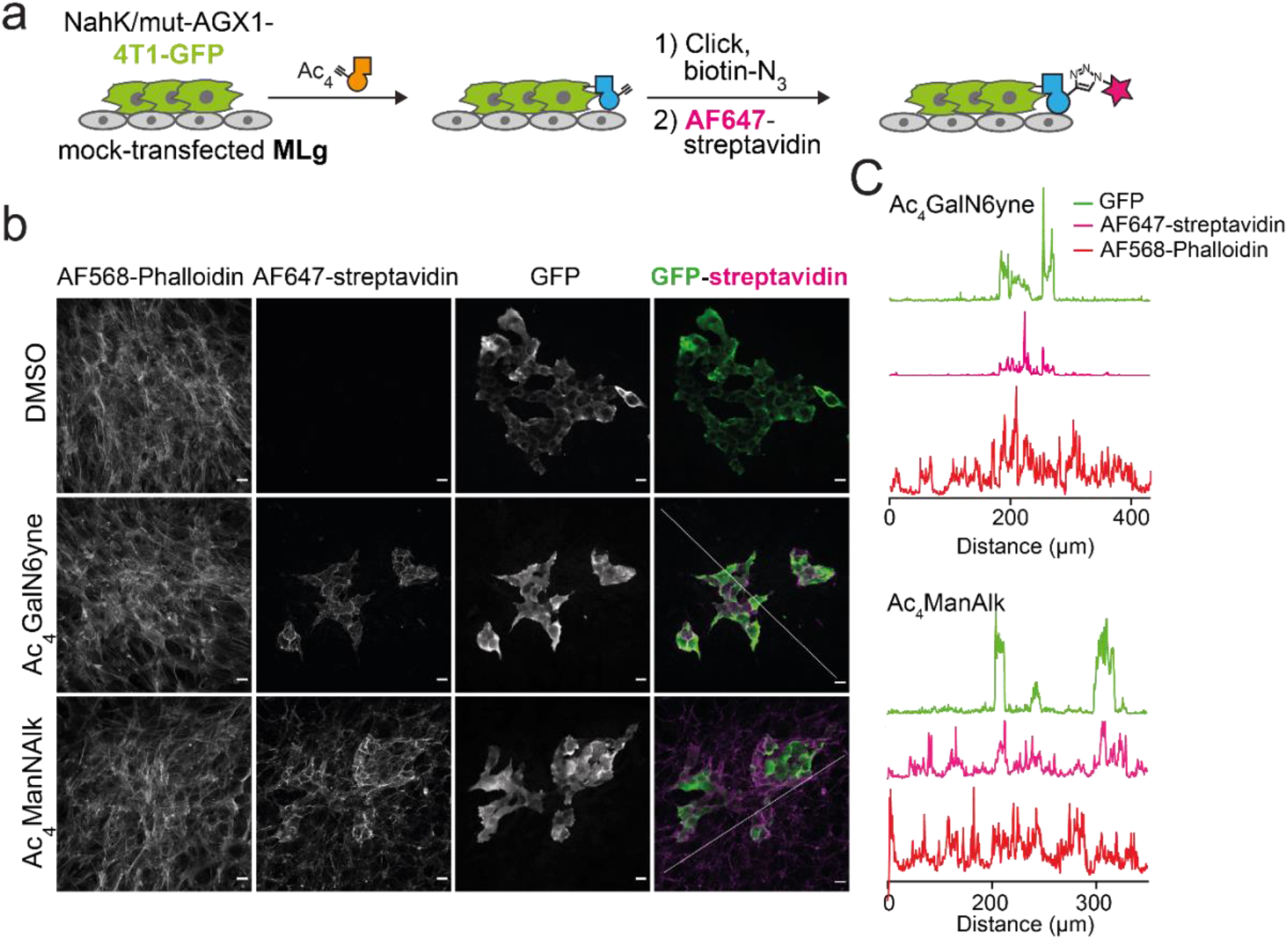
Bioorthogonal cell-specific glycoprotein tagging in co-culture. **a,** schematic of the 4T1-MLg co-culture experiment. Green fluorescent protein (GFP)-expressing 4T1 cells transfected with NahK/mut-AGX1 should be selectively positive for AF647-labelling in BOCTAG. **b,** fluorescence microscopy, using co-cultures fed with 50 μM Ac_4_GalN6yne or 50 μM Ac_4_ManNAlk as well as Alexafluor568-phalloidin as a counterstain. **c**, intensity profile of fluorescent signal between GFP and AF647 in Ac_4_GalN6yne- (top) or Ac_4_ManNAlk-fed (bottom) co-cultures. Scale bar, 20 μm. The intensity profile of GFP, AF647-Streptavidin and AF568-Phalloidin signals was measured along a diagonal line drawn along the fluorescent image. Data are representative of one out of two independent experiments.

### Assessing and manipulating the glycan types tagged by GalN6yne

We next sought to assess and expand the glycan subtypes targeted by our MOE approach. We were prompted by our recent findings that GalNAc analogues with bulky *N*-acyl chains such as GalN6yne are not incorporated into O-GalNAc glycans by WT-GalNAc-Ts (Fig. 4a).^23,27,34^ We have created GalNAc-T mutants termed BH-GalNAc-Ts (for “bump-and-hole engineering”, the process used to design the mutants) that selectively use such chemically tagged UDP-GalNAc analogues in glycosylation reactions.^23,27,34^ We stably co-expressed WT- or BH-versions of GalNAc-T1 or T2 from plasmids also encoding NahK and mut-AGX1 in K-562 cells (Fig. 4a). Expression of BH-GalNAc-Ts increased the intensity of in-gel fluorescence more than sevenfold over expression of WT-GalNAc-Ts when cells were fed with Ac_4_GalN6yne (Fig. 4b). WT-AGX1 expressing cells lacked UDP-GalN6yne/UDP-GlcN6yne biosynthesis and did not show any discernible fluorescent signal over vehicle control DMSO (Fig. 1c). We assessed the subtypes of the chemically tagged glycans by digestion with the hydrolytic enzymes PNGase F (reduces N-glycosylation), StcE (digests mucin-type glycoproteins) and OpeRATOR (digests O-GalNAc glycoproteins in the presence of the sialidase SialEXO) prior to in-gel fluorescence.^35^ In cells expressing NahK, mut-AGX1 and WT-GalNAc-Ts, fluorescent labelling was slightly sensitive to PNGase F treatment, indicating that the major target structures are N-glycoproteins in these cells (Fig. 4c). Co-expression of BH-GalNAc-Ts led to additional highly intense fluorescent signal of a small number of O-glycoproteins with sensitivity to both StcE and OpeRATOR/SialEXO (Fig. 4c). Thus, BH-GalNAc-Ts broaden the target scope of chemical tagging to include O-GalNAc glycoproteins with high incorporation efficiency. Concomitant with this finding, we performed quantitative MS-proteome analysis by SILAC of cell lines expressing NahK/mut-AGX1/BH-GalNAc-T2 (BH-T2). In contrast to cells expressing only NahK/mut-AGX1/WT-T2 (Fig. 2c), we observed an increase from 37% to 61% of O-GalNAc glycoproteins^31–33^ in the enriched protein fraction (Supplementary Table1).

**Fig. 4:**
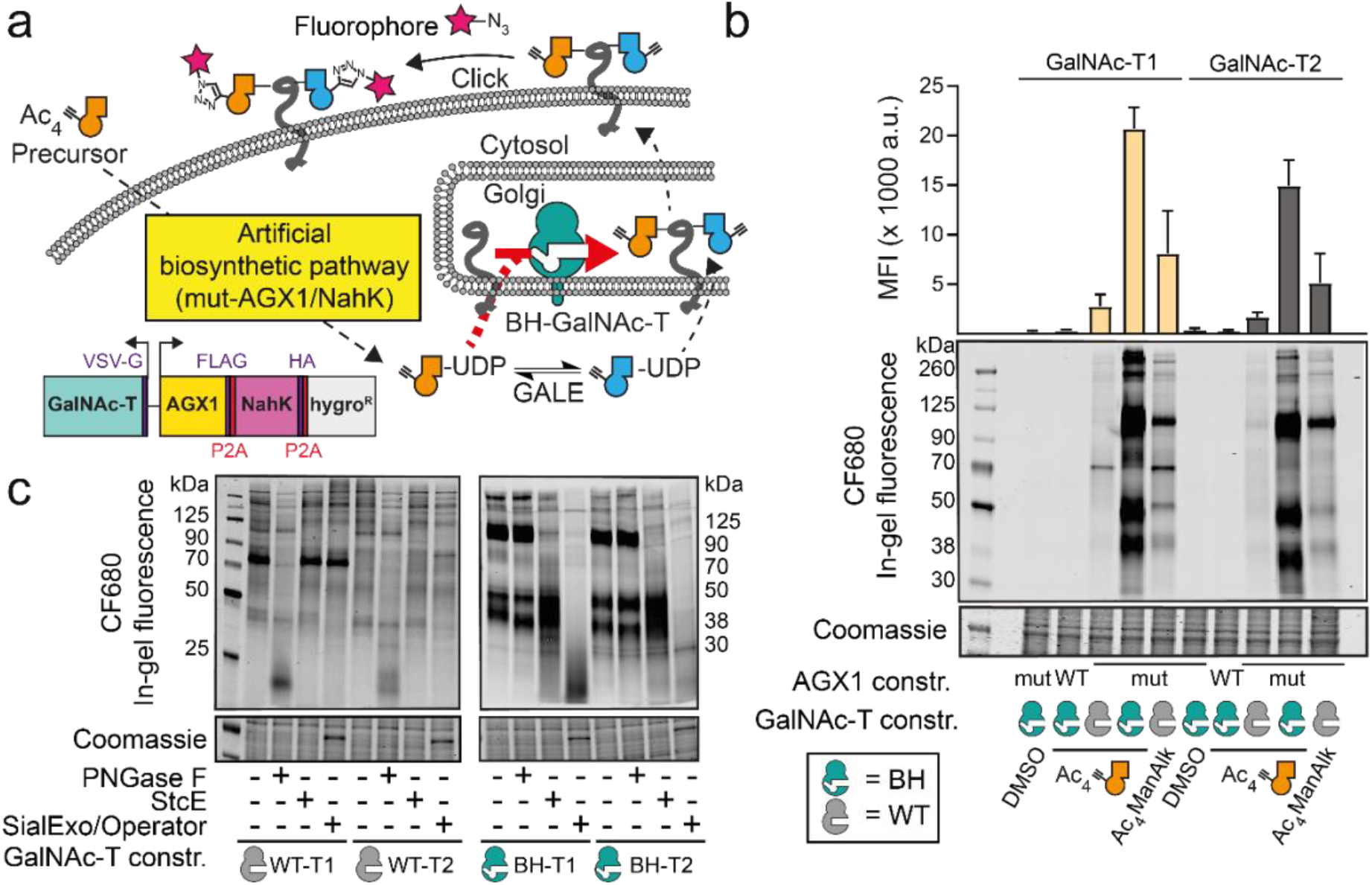
Enhancement of programmable glycoprotein tagging by expression of BH-GalNAc-Ts. **a,** strategy of expanding glycoprotein tagging to include O-GalNAc glycans. Expression of BH-GalNAc-Ts selectively engineered to accommodate bulky chemical tags enhances O-GalNAc tagging in cells expressing NahK/mut-AGX1. **b,** evaluation of tagging efficiency by feeding transfected K-562 cells DMSO, 1 μM Ac_4_GalN6yne or 2 μM Ac_4_ManNAlk. Tagging was analysed by in-gel fluorescence and quantification by densitometry as means + SD from three independent experiments. **c,** assessment of tagged glycan subtypes by treating the samples of cells fed with Ac_4_GalN6yne analysed in **b** with hydrolytic enzymes. Data are representative of one out of a total of four replicate labelling experiments performed on two different days.

### MS-based validation of cell-type specific labelling in co-culture models

We then validated BOCTAG as a strategy for cell-specific MS-glycoproteome analysis. We chose a co-culture model between murine 4T1 and human MCF7 breast cancer cell lines, opting to distinguish labelled glycoproteins with species-specific peptide sequences by label-free quantitative (LFQ) LC/MS-MS analysis. We transfected cells with either NahK/mut-AGX1/BH-GalNAc-T2 (termed “BOCTAG-T2”) or empty plasmid (pSBbi-Hyg, mock), co-cultured murine and human cells overnight and subsequently fed the co-cultures with either Ac_4_GalN6yne or vehicle DMSO. Chemically tagged glycoproteins in the secretome were reacted with acid-cleavable biotin-picolyl azide by CuAAC and enriched on neutravidin magnetic beads (Fig. 5a). On-bead digest yielded a peptide fraction and left glycopeptides bound to beads to be separately eluted with formic acid^24,36,37^. Peptide samples were assessed by LFQ MS-proteomics in two independent experiments, choosing a 8-fold enrichment and a p-value of 0.1 as cut-offs. We observed species-specific protein enrichment: BOCTAG-T2-expressing 4T1 cells led to 132 selectively enriched murine peptides while BOCTAG-T2-expressing MCF7 cells allowed detection of 24 selectively enriched human peptides when co-cultured with mock-transfected cells of the respective other species (Fig. 5b, Supplementary Table2). Only two human peptides and one murine peptide were found in the enriched datasets from the corresponding other species.

**Fig. 5:**
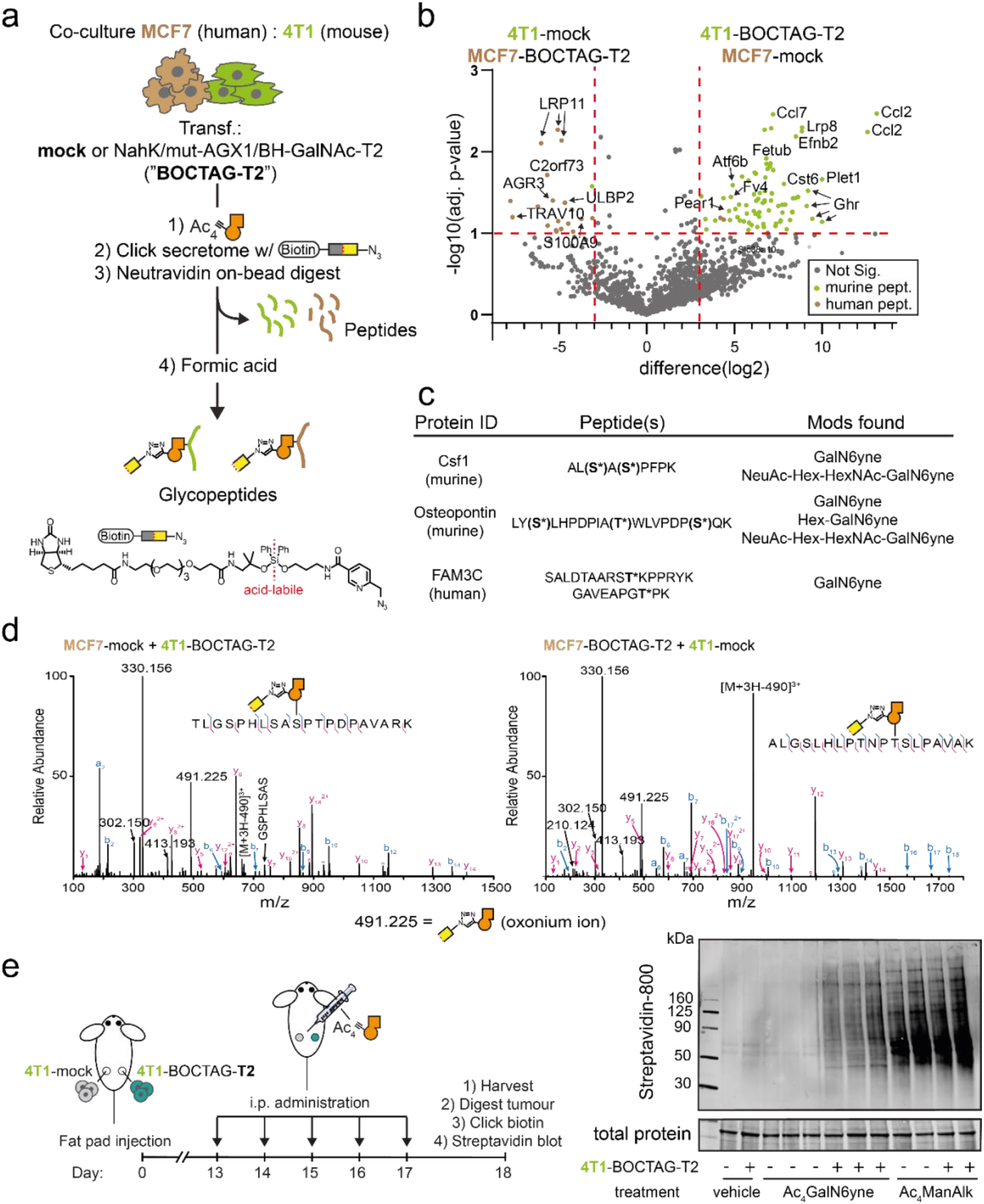
BOCTAG labels glycoproteins in a cell-specific manner in co-culture and *in vivo*. **a,** cell-selective enrichment and MS-glycoproteome analysis of murine-human co-culture systems. MCF7 and 4T1 cells transfected as indicated were co-cultured overnight and treated with DMSO or 10 μM Ac_4_GalN6yne for 24h. Secretome was subjected to CuAAC with acid-cleavable biotin-picolyl azide and enriched on neutravidin beads. On-bead digest yielded peptide fractions while acid treatment of beads yielded glycopeptide fractions. **b,** MS analysis of peptide fractions from **a** by choosing 8-fold enrichment and a p-value of 0.1 as cut-offs. Species-specific peptides are indicated. Data are from two independent experiments. **c,** examples of enriched glycopeptides and glycoforms. Asterisk annotates glycosylation sites; parentheses indicate potential glycosylation sites that could not be confidently assigned. **d,** HCD spectra of homologous glycopeptides from murine (left) and human (right) origins. Peptide sequences were confirmed by ETD (Fig S9c). **e,** *in vivo* glycoproteome tagging by BOCTAG-T2. Tumours were grown in fat pads of mice as described. BOCTAG-T2 and mock tumours were grown in the same mouse treated systemically by intraperitoneal (i.p. administration) for five days with 300 mg/kg Ac_4_GalN6yne, Ac_4_ManNAlk or the corresponding volume of vehicle. Tumours were harvested, lysed, subjected to CuAAC with biotin-picolyl azide and analysed by streptavidin blot. Hex = Hexose, e.g. galactose; NeuAc = *N*-acetylneuraminic acid; HexNAc =*N*-acetylhexosamine, e.g. GlcNAc. mock: pSBbi-Hyg.

BOCTAG-T2 allows for cell-specific glycosylation site identification. Using a tandem MS technique consisting of higher energy collision dissociation (HCD)-triggered electron transfer dissociation (ETD), we identified 37 specific glycosylation sites on 57 murine glycopeptides from 4T1 cells and 9 specific glycosylation sites on 12 human glycopeptides from MCF7 cells in secretome samples (Supplementary Table2). Our data indicated glycosylation of homologous glycopeptides from murine and human origins in pro-X carboxypeptidase in secretome (Fig. 5d, fig. S9c). We also performed an MS-glycoproteomics experiment in lysate from the 4T1/MCF7 co-culture expressing BOCTAG-T2 or empty plasmid. We annotated a total of 4 specific glycosylation sites on 11 murine glycopeptides from 4T1 samples and 2 specific glycosylation sites on 8 human glycopeptides from MCF7 cells (Supplementary Table 3). Particularly, we identified a homologous glycopeptide from both human and murine glucosidase 2 (Fig. S9b). The presence of the chemical tag facilitated manual annotation of mass spectra in all cases due to the specific mass shift associated with the chemical modification, in line with our own previous results.^38^

### Bioorthogonal cell-specific tagging of glycoproteins in vivo

We next investigated the applicability of our BOCTAG strategy in an *in vivo* tumour model. Tumours were grown in the fat pads of NOD-SCID IL2Rgnull (NSG) mice, consisting of 4T1 cells expressing GFP and either BOCTAG-T2 (one fat pad) or no additional transgene (empty plasmid, another fat pad). These mice were intraperitoneally injected with Ac_4_GalN6yne, vehicle or Ac_4_MAnNAlk once daily for five consecutive days (Fig. 5e). At the end of the treatment, the tumours were harvested, homogenized, treated with biotin-picolyl azide under CuAAC conditions and the labelling analysed by streptavidin blot. A strong fluorescent signal was observed in the BOCTAG-T2 tumours treated with Ac_4_GalN6yne (Fig 5e). In contrast, tumours transfected with empty plasmid showed minimal labelling signal with either vehicle or Ac_4_GalN6yne treatment. All samples treated with Ac_4_ManNAlk irrespective of the presence of NahK/mut-AGX1/BH-T2 displayed strong fluorescent signal. These data demonstrated that glycoproteins are selectively tagged when NahK/mut-AGX1/BH-T2 are expressed in the tumour. We also performed intratumoral injections of either Ac_4_GalN6yne or DMSO and observed the same BOCTAG-T2-dependent labelling (Fig. S10a).

To evaluate the protein expression levels of NahK/mut-AGX1/BH-T2 *ex vivo*, part of the tumours were digested, plated and cells cultured. Protein expression of NahK/mut-AGX1/BH-T2 was assessed by Western blot and found to be comparable to expression levels before *in vivo* injection (Fig. S10b). Cells also generally retained the ability to incorporate Ac4GalN6yne-dependent chemical glycoproteome tagging (Fig. S10b).

## DISCUSSION

We developed BOCTAG to address two major shortcomings in prominent research fields such as cancer biology. First, there is still an unmet need for characterising proteins produced by a particular cell type. Glycans are a means to an end in this respect, and the large signal-to-noise ratio in our fluorescent labelling experiments indicates that BOCTAG allows for efficient protein tagging. The approach is complementary to other techniques, including the use of unnatural amino acids and proximity biotinylation.^39,40^ Second, directly incorporating glycans in the analysis will give insight into cell-type-specific glycosylation sites and glycan structures to add another dimension to proteome profiling. The presence of a modification that can be observed by MS and is a direct corollary of using chemical tools allows for further validation of enriched glycoproteins, facilitating glycoproteome analysis even in complex co-culture or *in vivo* settings. An artificial biosynthetic pathway was essential to ensure minimal background labelling while being able to supply the tagged sugar as an easy-to-synthesise MOE reagent. To this end, the use of the kinase NahK allows for use of a per-acetylated bioorthogonal sugar that is fundamental to *in vivo* use and in marked difference to highly unstable caged sugar-1-phosphates used previously.^19,27^ To enable BOCTAG, cells require transfection with at least two transgenes. However, the design of a multicistronic, transposase-responsive plasmid ensures that transfection efforts are straightforward.^41,42^ BOCTAG allowed us to selectively tag tumour glycoproteomes *in vivo*, highlighting the robustness of the approach. MOE reagents have been chemically caged to be released by enzymes overexpressed in cancer^43–45^.While independent of transfection, such targeting can be accompanied by substantial background labelling in non-cancerous tissue. BOCTAG allows for programmable glycoprotein tagging with remarkable signal-to-noise ratio, and an enabling technology that will transform our understanding of tumour-host interactions particularly in the context of protein glycosylation.

## Data availability

Mass spectrometry raw sequencing data will be made available in a public repository.

## ACKNOWLEDGEMENTS

We thank Mahmoud-Reza Rafiee for help with data analysis, Acely Garza-Garcia, Holly Douglas and Christelle Soudy for help with chromatography, and Kayvon Pedram for providing StcE. We thank Junwon Choi for UDP-sugar standards for chromatography. We thank Phil Walker for advice on vector choice, and Rocco D’Antuono of the Crick Advanced Light Microscopy STP for support and assistance in this work. We thank Phil East of the Bioinformatics and Biostatistics Science Technology Platform for help with data analysis. We are grateful for support by the Francis Crick Institute Cell Services and Peptide Chemistry Science Technology Platforms. This work was supported by the Francis Crick Institute (A. C., B. C., T. R., V. B., A. M., G. B.-T., H. F., Z. L., O. Y. T., C. R., P. S.-B., S. K., I. M., B. S.) which receives its core funding from Cancer Research UK (FC001749, FC001045), the UK Medical Research Council (FC001749, FC001045) and Wellcome Trust (FC001749, FC001045). This work was supported by the ERC (788231 to S. L. F.), the Wellcome Trust (218304/Z/19/Z to A. M. and B. S.). the EPSRC (EP/S013741/1 to T. K. and M. A. F., and EP/S005226/1 to S. L. F.), the BBSRC (BB/T01279X/1 to Z. L. and B. S., BB/M027791/1 and BB/M028836/1 to S. L. F., and BB/M02847X/1 to T. K. and M. A. F.) and the NIH (R01 CA200423 to C.R.B.). B.C. was supported by a Crick-HEI studentship funded by the Department of Chemistry at Imperial College London and the Francis Crick Institute. For the purpose of Open Access, the author has applied a CC BY public copyright licence to any Author Accepted Manuscript version arising from this submission.

