## Supporting Information for "Cell-specific Bioorthogonal Tagging of Glycoproteins"

Supporting Figures
Experimentals

Anna Cioce[a,b], Beatriz Calle[a,b], Tatiana Rizou[c,$], Sarah C. Lowery[d,$], Victoria Bridgeman[c,$], Keira E. Mahoney[d,$], Andrea Marchesi[a,b], Ganka Bineva-Todd[b], Helen Flynn[e], Zhen Li[a,b], Omur Y. Tastan[b], Chloe Roustan[f], Pablo Soro-Barrio[g], Thomas M. Wood[h,k], Tessa Keenan[i], Peter Both[j], Kun Huang[j,l], Fabio Parmeggiani[j,m], Ambrosius P. Snijders[e], Mark Skehel[e], Svend Kjaer[f], Martin A. Fascione[i], Carolyn R. Bertozzi[h], Sabine Flitsch[j], Stacy A. Malaker[d], Ilaria Malanchi[c], Benjamin Schumann[a,b,*]

[a]Department of Chemistry, Imperial College London, 80 Wood Lane, W12 0BZ, London, United Kingdom.
[b]Chemical Glycobiology Laboratory, The Francis Crick Institute, 1 Midland Rd, NW1 1AT London, United Kingdom.
[c]Tumour-Host Interaction Laboratory, The Francis Crick Institute, 1 Midland Rd, NW1 1AT London, United Kingdom.
[d]Department of Chemistry, Yale University, 275 Prospect Street, New Haven, CT 06511, United States.
[e]Proteomics Science Technology Platform, The Francis Crick Institute, NW1 1AT London, United Kingdom.
[f]Structural Biology Science Technology Platform, The Francis Crick Institute, NW1 1AT London, United Kingdom.
[g]Bioinformatics & Biostatistics Science Technology Platform, The Francis Crick Institute, NW1 1AT London, United Kingdom.
[h]Department of Chemistry, Stanford University, Stanford, CA 94305, USA.
[i]Department of Chemistry, University of York, YO10 5DD York, United Kingdom.
[j]Manchester School of Chemistry & Institute of Biotechnology, The University of Manchester, M1 7DN Manchester, United Kingdom.
[k]current address: Massachusetts Institute of Technology, Cambridge, USA
[l]current address: Department of Chemistry and Biochemistry, University of Maryland, Baltimore, MD 21250, USA.
[m]current address: Department of Chemistry, Materials and Chemical Engineering "G. Natta", Politecnico di Milano, 20131 Milano, Italy
[$]These authors contributed equally.

### Supporting Figures

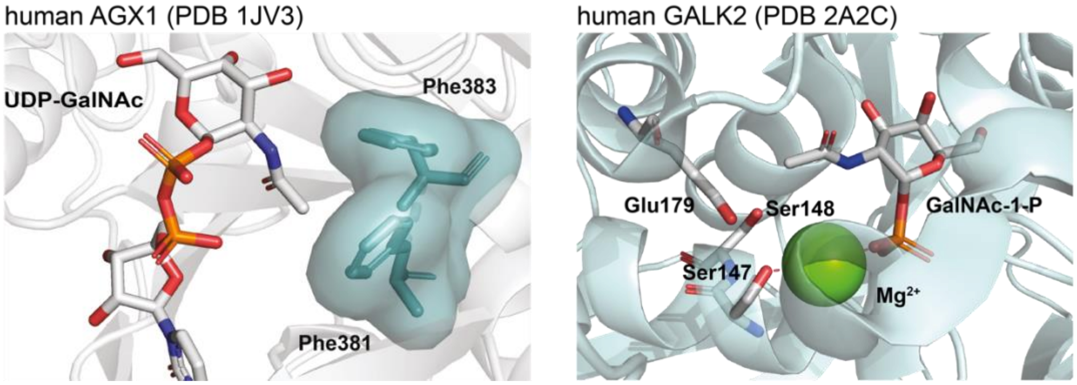


**Fig. S1: Active site architectures of human enzymes of the GalNAc salvage pathway.** In AGX1, the *N*-acyl side chain in UDP-GalNAc is in proximity to Phe381 and Phe383. In GALK2, the *N*-acyl side chain of GalNAc-1-phosphate is in proximity with amino acids forming a hydrogen network (Glu179, Ser147 and Ser148). Structures are modelled based on protein databank acquisition numbers 1JV3[1] and 2A2C.[2]

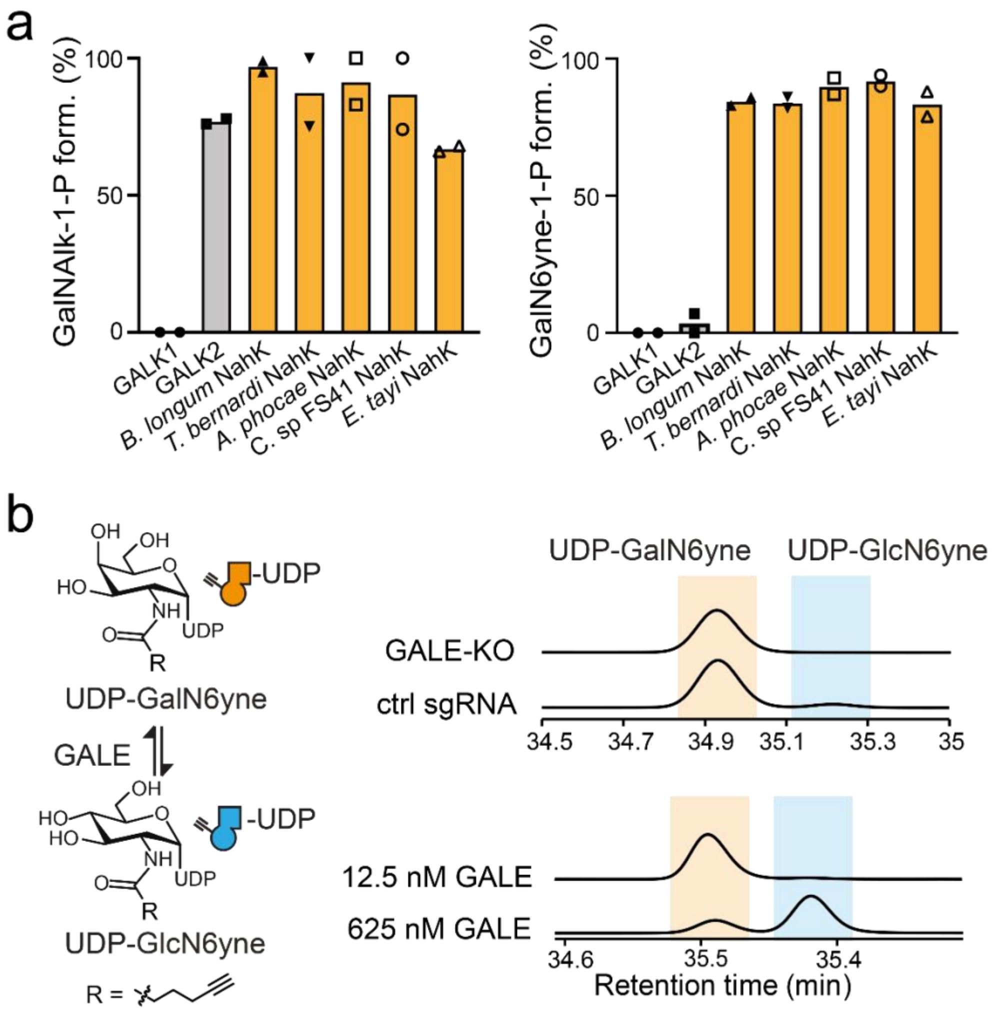


**Fig. S2: Evaluation of enzymatic turnover of GalN6yne-based metabolites. a,** *in vitro* GalN6yne-1-phosphate and GalNAlk-1-phosphate formation by human and bacterial GalNAc kinases, as assessed by LC-MS and integrated ion count. Data are individual data points and means from two individual experiments. **b,** ion pair HPLC traces of *in vitro* epimerisation assays, using UDP-GalN6yne as substrate and either cell lysates from control or GALE-KO K-562 cells or two different concentrations of purified GALE as enzyme sources.[3] Data are representative of two independent replicates (lysate samples) or from one experiment (isolated GALE samples). Traces depict relative intensity of absorbance at 260 nm.

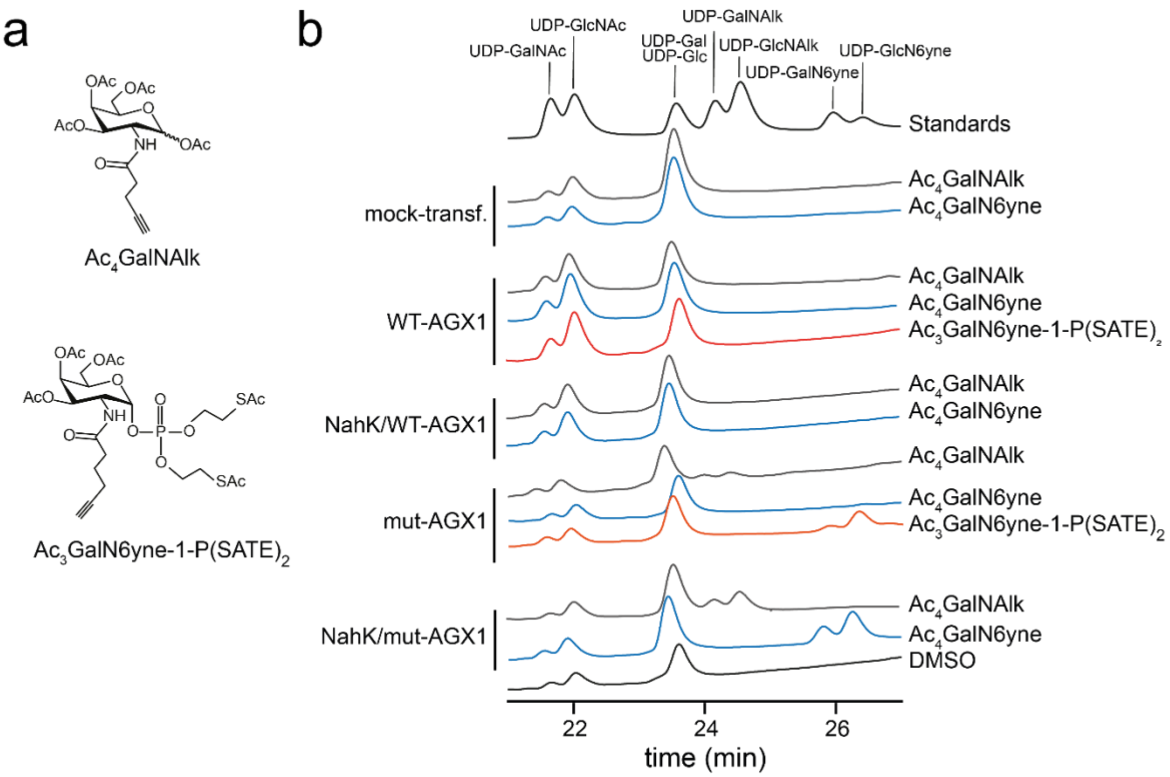


**Fig. S3: Biosynthesis of chemically tagged UDP-sugars by metabolic engineering. a,** structures of two MOE reagents used herein. $Ac_3GalN6yne$-1-P(SATE)$_2$ is a caged precursor of GalN6yne-1-phosphate. **b,** biosynthesis of UDP-sugars in cells stably transfected with metabolic enzymes, as assessed by high performance anion exchange chromatography (HPAEC). Chromatograms were normalized on an external standard. Synthetic UDP-sugars served as standards. Data are from one representative out of two independent experiments performed on two different days.

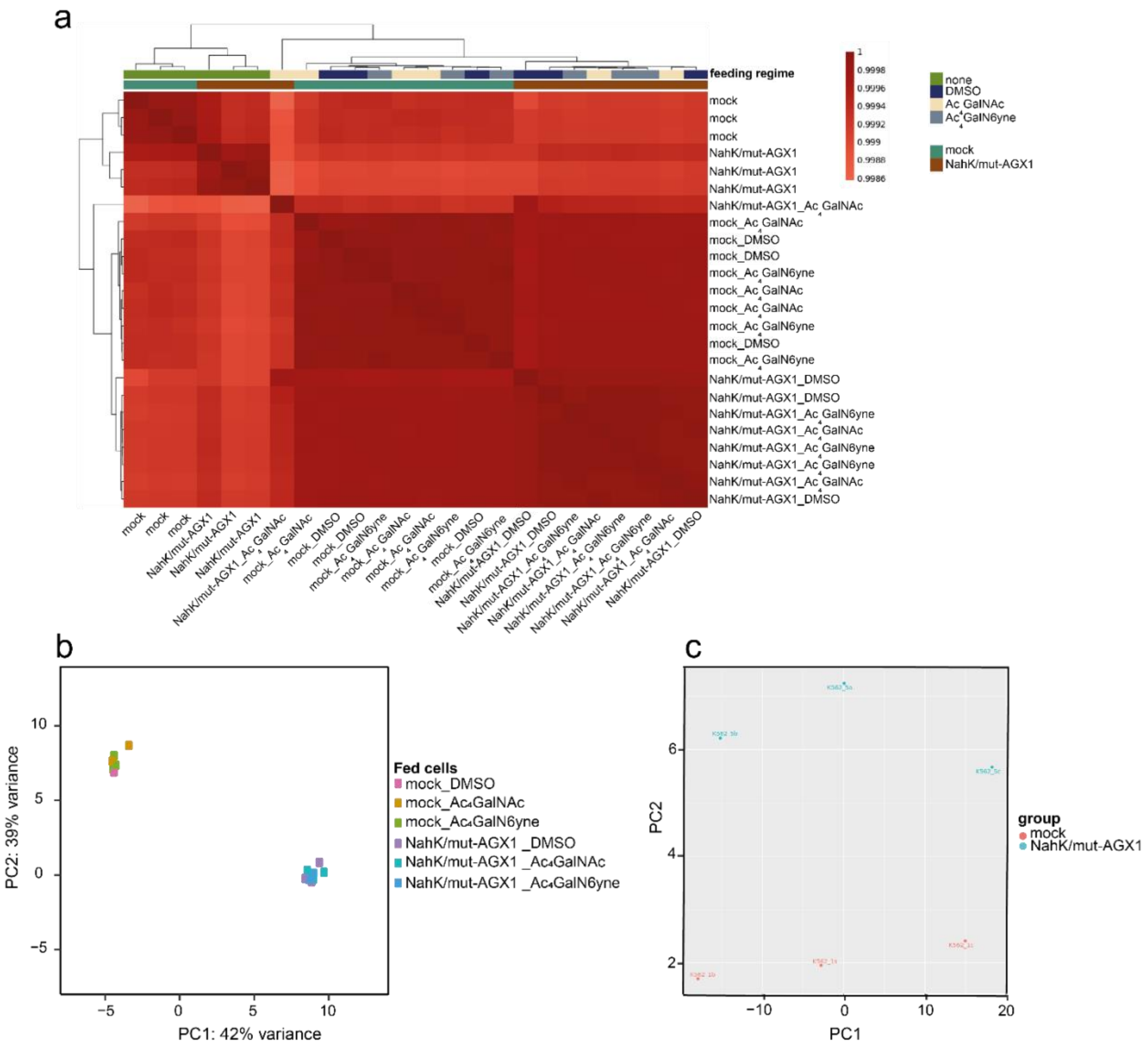


**Fig. S4: transcriptomics analysis of K-562 cells at different transfection conditions and feeding regimes.** RNAs were extracted from K-562 cells transfected with either pSBbi-GH (mock) and pSBbi- NahK/mut-AGX1[4] at different feeding regimes (either unfed, DMSO, 10 µM Ac$_4$GalNAc or 10 µM Ac$_4$GalN6yne). **a,** Correlation plot and **b,** Principal component analysis (PCA) plot of transfected K-562 cells at different feeding regimes. Data are from one representative out of a total of three replicates collected the same day. **c,** Previous experiment performed on RNA extracted from K-562 cells, transfected with either pSBbi-GH (mock) and pSBbi- NahK/mut-AGX1, showed significant variation in PCA associated to the different day of collection.

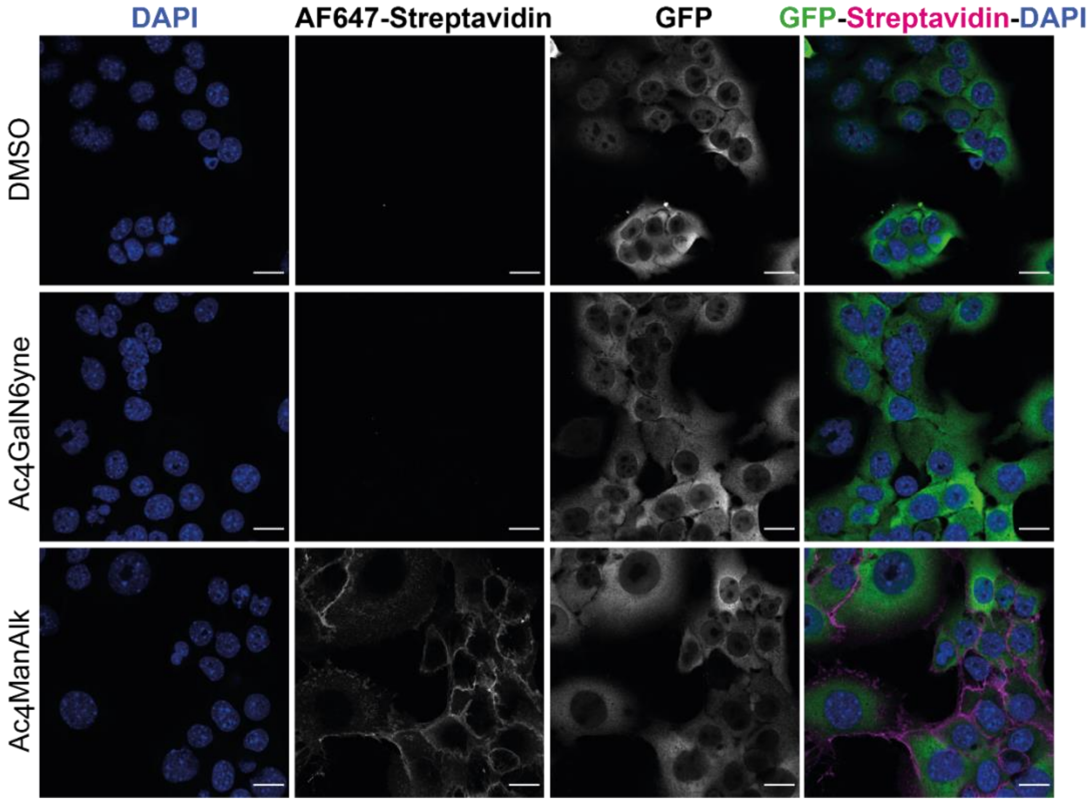


**Fig. S5:** Fluorescence microscopy of GFP-expressing 4T1 cells, transfected with empty plasmid pSBbi-Hyg, fed overnight with either DMSO, 50 µM $Ac_4GalN6yne$ or 50 µM $Ac_4ManAlk$, treated with biotin-picolyl azide under CuAAC conditions and visualized with Alexafluor647-Streptavidin. Scale bar, 20 µm.

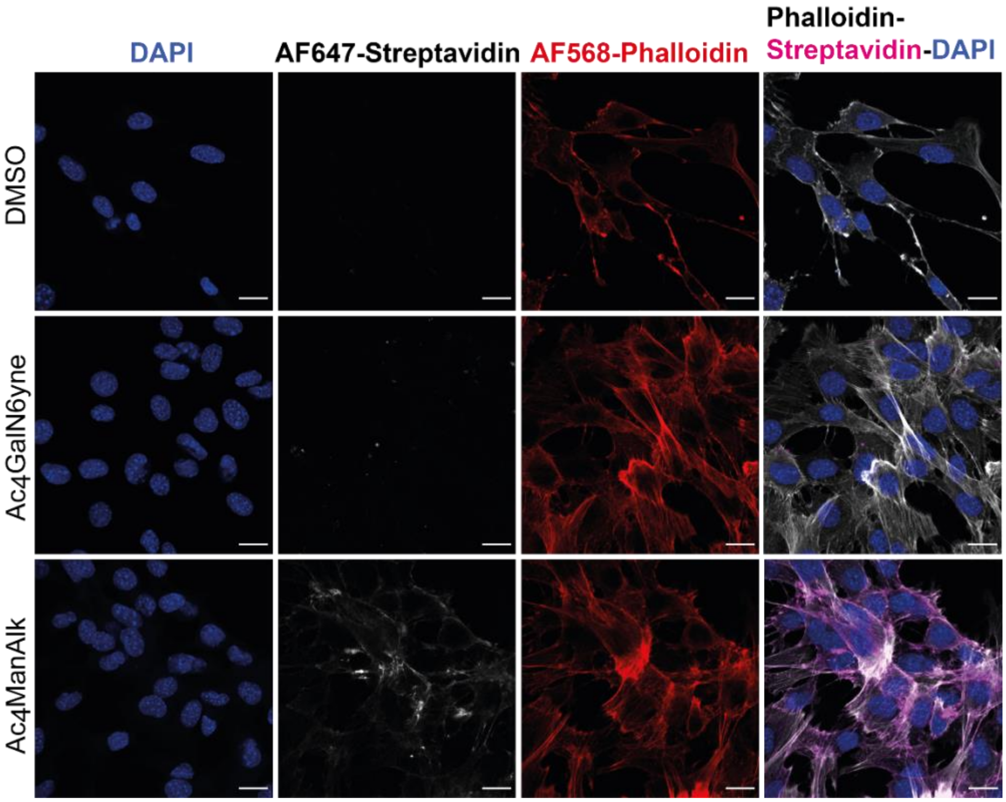


**Fig. S6:** Fluorescence microscopy of non-transfected MLg cells fed overnight with DMSO, 50 µM $Ac_4$GalN6yne or 50 µM $Ac_4$ManAlk treated with biotin picolyl azide under CuAAC conditions and visualized with Alexafluor647-Streptavidin. Scale bar, 20 µm.

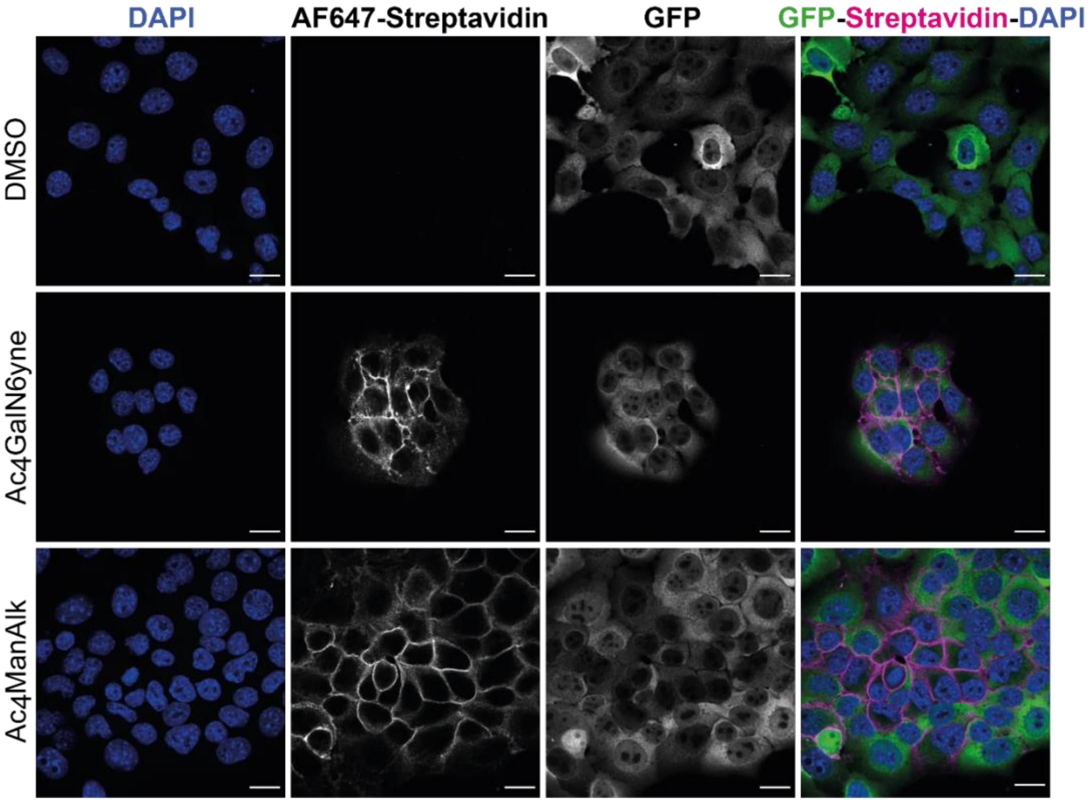


**Fig. S7:** Fluorescence microscopy of GFP-expressing 4T1 cells, transfected with pSBbi- NahK/mut-AGX1, fed overnight with either 50 µM $Ac_4$GalN6yne or 50 µM $Ac_4$ManAlk treated with biotin-picolyl azide under CuAAC conditions and visualized with Alexafluor647-Streptavidin. Scale bar, 20 µm.

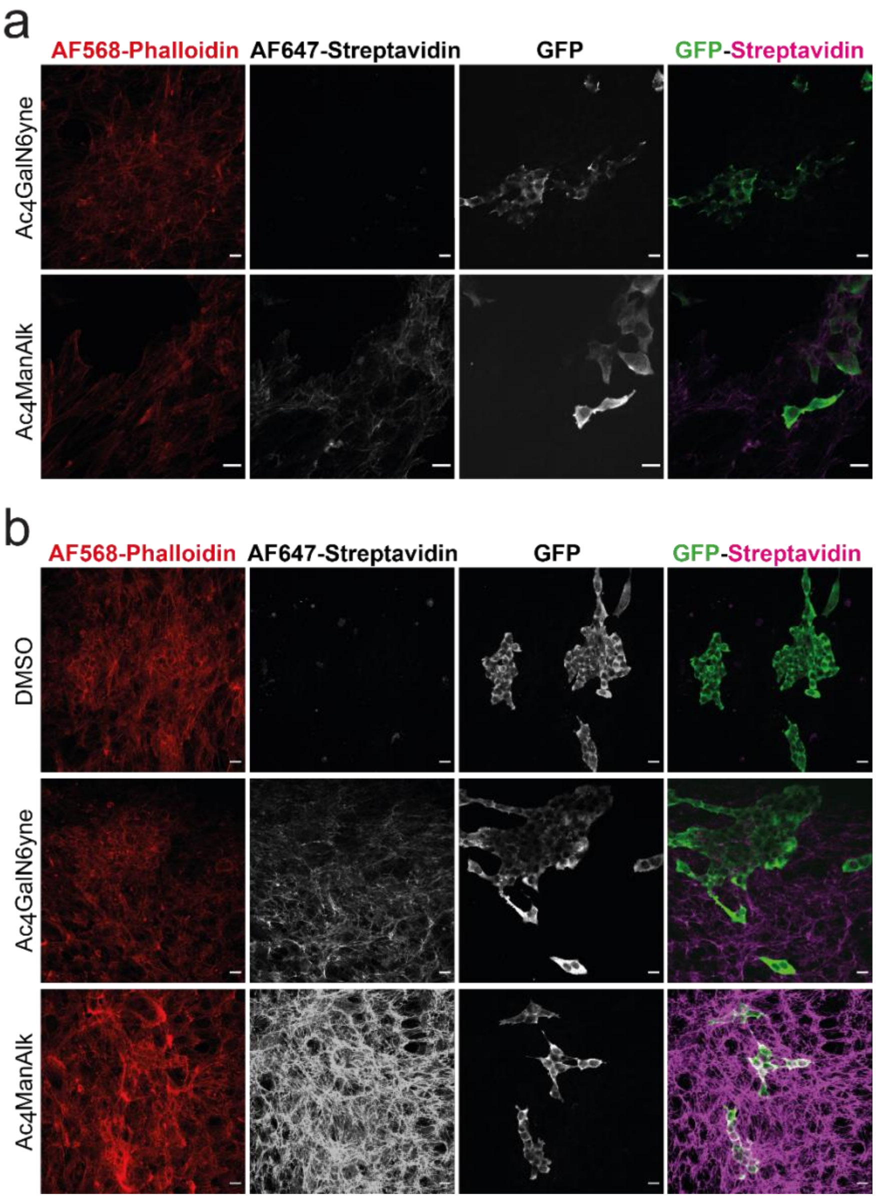


**Fig. S8:** Maximum intensity projection from a z-stack acquisition of: **a,** GFP-expressing 4T1 and MLg cells, both transfected with pSBbi-Hyg empty plasmid, in a co-culture system fed overnight with either 50 µM $Ac_4$GalN6yne or 50 µM $Ac_4$ManAlk. **b,** GFP-expressing 4T1 and MLg cells, both transfected with pSBbi-NahK/mut-AGX1 plasmid, in a co-culture system fed overnight with either DMSO, 50 µM $Ac_4$GalN6yne or 50 µM $Ac_4$ManAlk. Co-culture samples in both A and B are treated with biotin picolyl azide under CuAAC conditions and visualized by Alexafluor647-Streptavidin. Scale bar, 20 µM.

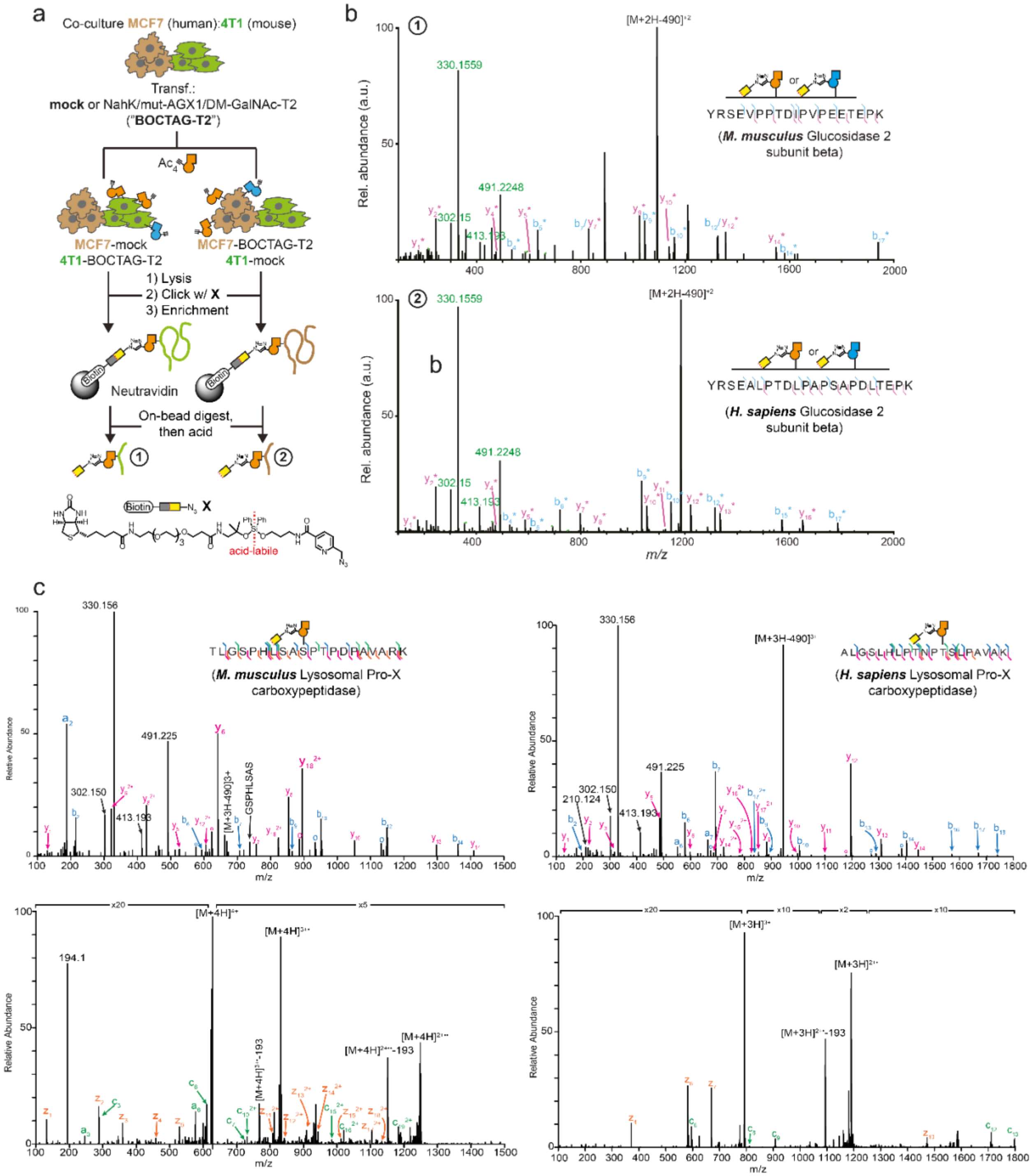


**Fig. S9: Proteomics and glycoproteomics analysis of lysates of co-cultured human and murine samples.** **a,** cell-selective enrichment and MS-glycoproteomics of murine-human co-culture systems. MCF7 and 4T1 cells transfected as indicated were co-cultured overnight and treated with DMSO or 10 µM $Ac_4$GalN6yne for 24h. Lysate was subjected to CuAAC with acid-cleavable biotin picolyl azide **X** and enriched on neutravidin beads. On-bead digest yielded peptide fractions while acid treatment of beads yielded glycopeptide fractions. **b,** HCD (Higher-energy collisional dissociation) spectra of homologous glycopeptides from murine (top) and human (bottom) origins. **c,** HCD and Electron Transfer Dissociation Mass Spectrometry (ETD) spectra of homologous glycopeptides from murine (left) and human (right) origins enriched in secretome.

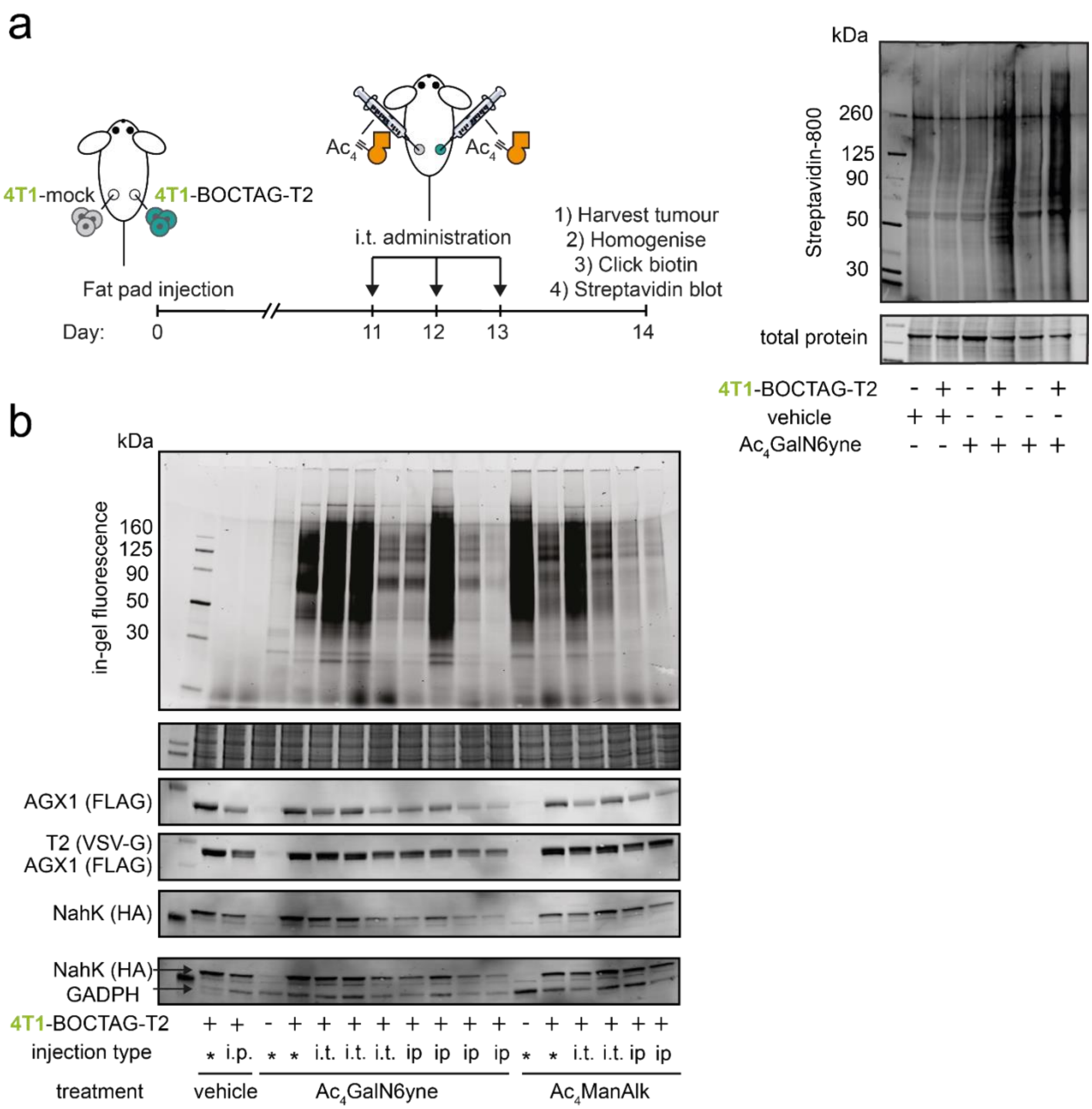


**Fig. S10: BOCTAG labels glycoproteins in a cell-specific manner *in vivo*. a,** BOCTAG-T2 and mock tumours were grown in the same mouse treated systemically for three days with 300 mg/kg $Ac_4GalN6yne$ or the corresponding volume of vehicle (5% (v/v) DMSO/ PEG-400) by intratumoral (i.t.) injection. Tumours were harvested, lysed, subjected to CuAAC with biotin-picolyl azide and analysed by streptavidin blot. **b,** after the experiment in **Fig.5** and **fig. S10a**, BOCTAG-T2 cells were collected from intratumoral (i.t.) and intraperitoneal (i.p.) treated tumours, plated for 10 days in DMEM growing media and fed with either DMSO, $Ac_4GalN6yne$ or $Ac_4ManNAlk$. Glycoprotein labelling efficiency of primary cells was validated by CuAAC with CF680-picolyl azide. Western blot was performed to assess levels of expression of BOCTAG-T2 enzymes. i.p.= intraperitoneal injection, i.t.= intratumoral injection, *= parental 4T1-mock and 4T1-BOCTAG-T2 cell lines.

### Experimentals

The compounds $Ac_4$ManNAlk and $Ac_4$GalNAlk were purchased from Click Chemistry Tools (Scottsdale, USA). GalNAlk, GalN6yne, GalNAlk-1-phosphate, $Ac_3$GalN6yne-1-P(SATE)$_2$ and Ac4GalN6yne were prepared as reported.[5–7]

Adobe Illustrator was used to assemble figures and to re-label plot axes.

*In vitro phosphorylation of GalN6yne*

NahK from *B. longum* was either recombinantly expressed[8] or purchased from Chemily (Peachtree Corners, USA). All other bacterial NahKs were produced by Prozomix Ltd. (Haltwhistle, UK). Human kinases GALK1 and GALK2 were expressed following a standard baculovirus plasmid transfer expression protocol in SF21 insect cells as described before,[9] and purified by GST affinity and size exclusion chromatography steps.

Reactions were run in 50 µL volume, containing GalN6yne (5 mM), Adenosine triphosphate (ATP, 10 mM), $MgCl_2$ (10 mM), Tris-HCl pH 8 (100 mM) and kinase (20 µg). Reactions were run for 4 h at 37 °C. Reactions were diluted with 50 µL methanol, cooled to -20 °C for 2 h and centrifuged (18000 g, 30min) to remove any precipitated enzyme. Supernatants were analysed by UPLC (Ultra Performance Liquid Chromatography)-MS equipped with ACQUITY UPLC BEH Glycan 1.7 µm 2.1x50 mm column (90-65% buffer B over 17 minutes; buffer A: 10 mM ammonium formate pH 4.5, buffer B: 10 mM ammonium formate 90/10 (v/v) acetonitrile/water). Estimated conversion was obtained by analysing a 5 mM product standard in the same analysis conditions as above, extracting product mass and comparing ion count with the ion count of extracted mass of product in the samples.

*In vitro synthesis of UDP-GalN6yne*

AGX1 constructs were expressed and purified as reported before.[9] Reactions were run in 15 µL volume, containing GalN6yne-1-phosphate (2.5 mM), $MgCl_2$ (5 mM), Tris-HCl pH 8 (75 mM), BSA (1 mM), Uridine triphosphate (UTP, 5 mM), pyrophosphatase (PmPpA, Chemily, 0.045 U), recombinant WT- or mut-AGX1 (125 nM). Reactions were run for either 2 h or 16 h at 37 °C. 7 µL of each reaction were diluted with 7 µL of acetonitrile, cooled on ice for 30 min and centrifuged (18000 g, 30 min) to remove any precipitated enzyme. Supernatants were analysed by UPLC-MS equipped with ACQUITY UPLC BEH Glycan 1.7 µm 2.1x50 mm column (90-65% buffer B over 17 minutes; buffer A: 10 mM ammonium formate pH 4.5, buffer B: 10 mM ammonium formate 90/10 (v/v) acetonitrile/water).

*Plasmids and Cell lines*

AGX1$^{WT}$ and AGX1$^{F383A}$ were introduced into the plasmid pSBbi using a previously reported cloning strategy.[3] The pSBbi plasmid was a gift from Eric Kowarz (Addgene plasmid #60514; http://n2t.net/addgene:60514; RRID:Addgene_60514).[4] pCMV(CAT)T7-SB100 was a gift from Zsuzsanna Izsvak (Addgene plasmid #34879; http://n2t.net/addgene:34879; RRID:Addgene_34879).[10] NahK from *Bifidobacterium longum* (E8MF12) was codon-optimised for human expression and inserted into pSBbi-AGX1$^{WT}$ and pSBbi-AGX1$^{F383A}$ by GeneArt (Thermo Fisher, Waltham, USA), containing 2A self-cleaving peptides and a C-terminal HA tag, to give the plasmids pSBbi-AGX1$^{WT}$-NahK, pSBbi-AGX1$^{F383A}$-NahK. WT-or "bump-and-hole" (BH) engineered versions of GalNAc-T1 or GalNAc-T2 were inserted into these plasmids using an SfiI cloning strategy according to Schumann et al.[6] to give the plasmids pSBbi-AGX1$^{WT}$-NahK-T1$^{WT}$, pSBbi-AGX1$^{WT}$-NahK-T1$^{I238A/L295A}$, pSBbi-AGX1$^{F383A}$-NahK-T1$^{WT}$, pSBbi-AGX1$^{F383A}$-NahK-T1$^{I238A/L295A}$, pSBbi-AGX1$^{WT}$-NahK-T2$^{WT}$, pSBbi-AGX1$^{WT}$-NahK-T2$^{I253A/L310A}$, pSBbi-AGX1$^{F383A}$-NahK-T2$^{WT}$, pSBbi-AGX1$^{F383A}$-NahK-T2$^{I253A/L310A}$. pSBbi-AGX1$^{WT}$ and pSBbi-AGX1$^{F383A}$ were used as template to prepare the following plasmids: pSBbi-AGX1$^{WT}$, pSBbi-AGX1$^{F383A}$, pSBbi-AGX1$^{WT}$-NahK, pSBbi-AGX1$^{F383A}$-NahK, pSBbi-AGX1$^{WT}$-NahK-T1$^{WT}$, pSBbi-AGX1$^{WT}$-NahK-T1$^{BH}$, pSBbi-AGX1$^{F383A}$-NahK-T1$^{WT}$, pSBbi-AGX1$^{F383A}$-NahK-T1$^{BH}$, pSBbi-AGX1$^{WT}$-NahK-T2$^{WT}$, pSBbi-AGX1$^{WT}$-NahK-T2$^{BH}$, pSBbi-AGX1$^{F383A}$-NahK-T2$^{WT}$, pSBbi-AGX1$^{F383A}$-NahK-T2$^{BH}$.

AGX1, NahK and GalNAc-T1/T2 constructs were tagged with C-terminal FLAG, HA and VSV-G tags, respectively. The full plasmid sequence of pSBbi-AGX1$^{F383A}$-NahK-T2$^{I253A/L310A}$ is provided in the Appendix (see below).

All cells were screened for contamination by mycoplasma and other cell lines by the Crick Cell Services Science Technology Platform. K-562 cells were propagated in RPMI (Thermo Fisher) with 10% (v/v) FBS, penicillin (100 U/mL) and streptomycin (100 µg/mL). 4T1(GFP-expressing) and murine MLg fibroblast cells were maintained in DMEM (Thermo Fisher) with 10% (v/v) FBS, penicillin (100 U/mL) and streptomycin (100 µg/mL).

K-562 and 4T1(GFP) stably transfected with pSBbi-AGX1$^{WT}$ or pSBbi-AGX1$^{F383A}$ have been prepared previously.[6] K-562 were stably transfected with pSBbi-AGX1-NahK or pSBbi-AGX1-NahK-T1/T2 constructs or empty pSBbi-GH using Lipofectamine LTX (Thermo Fisher) according to the manufacturer's instructions, with a 20:1 (w/w) mixture of pSBbi and pCMV(CAT)T7-SB100 plasmid DNA. After 24 h, cells were harvested and selected in growth medium containing 150 µg/mL hygromycin B (Thermo Fisher) for 7-10 days to obtain stable cells._4T1 (GFP) and MLg were stably transfected with either pSBbi-AGX1$^{F383A}$-NahK or empty pSBbi-Hyg using Lipofectamine 3000 (Thermo Fisher) according to the manufacturer's specifications, with a 20:1 (w/w) mixture of pSBbi and pCMV(CAT)T7-pSB100 plasmid DNA. After 24 h, cells were harvested and selected in growth medium containing 100 µg/mL hygromycin B (Thermo Fisher) for 7-10 days to obtain stable cells.

*Analysis of nucleotide-sugar biosynthesis by High Performance Anion Exchange Chromatography*

Five million K-562 cells stably transfected with pSBbi-GH, pSBbi-AGX1$^{WT}$, pSBbi-AGX1$^{F383A}$, pSBbi-AGX1$^{WT}$-NahK and pSBbi-AGX1$^{F383A}$-NahK were fed with 100 µM (from a 100 mM stock solution in DMSO) membrane permeable precursor $Ac_4$GalN6yne, $Ac_4$GalNAlk, $Ac_3$GalN6yne-1-P(SATE)$_2$ or DMSO. After 16 h, cells were harvested, centrifuged at 500 *g*, 5 min, 4 °C and resuspended in PBS (1 mL). Cell pellets were resuspended in PBS (1 mL). 0.9 mL cell suspension transferred to O-ring tubes (1.5 mL, Thermo Fisher) and harvested. Zirconia/silica beads (0.1 mm, BioSpec, Bertlesville, USA) were added at a volume comparable to the cell pellet volume, followed by 1:1 acetonitrile/water (1 mL). Cells were lysed using a bead beater (FastPrep-24, MP Biomedicals, Santa Ana, USA) at 6 m/s for 30 s, and the cell lysate was cooled at 4 °C for 10 min. Samples were centrifuged (14000 g, 10 min, 4 °C), and the supernatant was transferred to a fresh tube. The solvent was evaporated by SpeedVac, and the residue was dissolved in LCMS-grade water (Thermo Fisher, 0.2-0.4 mL) containing 15 µM ADP-α-D-glucose (Sigma-Aldrich, St. Louis, USA). The solution was passed through a centrifuge filter (30 min, 14000 g) using a 3 kDa MWCO Amicon Ultra Centrifugal Filter Unit (Merck). The flow-through was evaporated by SpeedVac and the residue resuspended in 60µL of MQ water. High Performance Anion Exchange Chromatography (HPAEC) was used to analyse lysates.

HPAEC was carried out using an ICS-6000 equipped with a quaternary pump and a conductivity detector (data collection rate 5.0 Hz, cell temperature 35 °C) on an AS11 2x250 mm column and a 2x50 mm guard column (Thermo Fisher). Solvents were: A = water; B = 1 M NaOH. The gradient profile was as follows: 0 min 99.9% A, 0.1% B; 3 min 99.9% A, 0.1% B; 8 min 96.9% A, 3.1% B; 13 min 96.4% A, 3.6% B; 38 min 93% A, 7% B; 39 min 90% A, 10% B; 43 min 90% A, 10% B; 48 min 90% A, 10% B. Commercial or synthetic UDP-sugar standards (200-500 µM) were used as controls.

*Metabolic cell surface labelling and in-gel fluorescence*

K-562 cells stably transfected with pSBbi-based plasmids were seeded at a density of 250,000 cells/mL into well plates in growth medium without hygromycin. Cells were treated with $Ac_4$GalN6yne or $Ac_4$ManNAlk at the indicated concentrations or a corresponding volume of DMSO. Cells were grown for 20 h. Cells were harvested (500 g, 5 min, 4 °C) in a V-shaped 96 well plate and washed twice with 2% (v/v) FBS in PBS (Labelling Buffer, 0.2 mL). Cells were resuspended in Labelling Buffer (35 µL), treated with a solution of 200 µM $CuSO_4$, 1.2 mM BTTAA (Click Chemistry Tools, Scottsdale, USA), 5 mM sodium ascorbate, 5 mM aminoguanidinium chloride and 200 µM CF680 picolyl azide (Biotium) in Labelling Buffer (35 µL), and incubated for 7 min at room temperature on an orbital shaker. The click reaction was quenched with 3 mM bathocuproinedisulfonic acid (BCS) in PBS (35 µL). Cells were centrifuged, washed twice with Labelling Buffer and then with PBS, and treated with 100 µL of ice-cold Lysis Buffer (50 mM Tris-HCl pH 8, 150 mM NaCl, 1% (v/v) Triton X-100, 0.5% (v/v) sodium deoxycholate, 0.1% (w/v) SDS (Sodium dodecyl sulfate), 1 mM $MgCl_2$, and 100 mU/µL benzonase (Merck)) containing halt protease inhibitors. Cells were lysed for 20 min at 4 °C on an orbital shaker and centrifuged (1500 g, 20 min, 4 °C). Supernatant was transferred to a new plate and Pierce™ BCA Protein Assay kit (Thermo Fisher) was used to measure protein concentration. Loading buffer (a 1:1:1:0.5 (v/v/v/v) mixture of 1 M Tris-HCl pH 6.5, 80% (v/v) glycerol, 10% (w/v) SDS and 1 M dithiothreitol (DTT) was added; samples were run on a 10% or 4-20% Criterion™ gel (Bio-Rad, Hercules, USA) for SDS-PAGE, and imaged on an Odyssey CLx imager (LI-COR Biosciences, Lincoln, USA). For protein quantification by densitometry, the signal in each lane was integrated in the 700 nm channel using Image Studio Pro software (LI-COR Biosciences, Lincoln, USA). Total protein was stained with Coomassie using Acquastain (Bulldog Bio, Portsmouth, USA). Protein expression was assessed by Western blot with a different set of samples, using antibodies against FLAG tag (rabbit anti-FLAG antibody,1:1000, PA1-984B, invitrogen, , Carlsbad, USA), HA tag (rabbit anti-HA antibody, 1:1000, ab9110, abcam), VSV-G tag (goat anti-VSV-G, 1:2000, ab3861, abcam) and GADPH (rabbit anti-GAPDH, 1:5000, ab181602, abcam).

For enzymatic treatment, lysates from cell surface-labelled cells (5 µg protein) were diluted to 20 µL with 50 mM Tris-HCl pH 7.5 and 150 mM NaCl. Samples were either left untreated, or treated with 2 µL of a 1:10 dilution in PBS of commercial PNGase F (Promega, Madison, USA), 2 µL StcE (96.5 µg/mL solution in PBS)[11] or 2 µL of a mixture of SialEXO and OpeRATOR (4 U/µL in PBS, Genovis, Lund, Sweden) Samples were incubated for 2 h at 37 °C, briefly heated to 95 °C and cooled on ice. SDS-PAGE and in-gel fluorescence were performed as described above.

In-gel fluorescence and Western Blot images were visualized and processed on ImageStudioLite software (LI-COR Biosciences, Lincoln, USA) and cropped by Illustrator (Adobe, San Jose, USA).

*Transcriptomie analysis*

K-562 cells ($1x10^6$), transfected with pSBbi-GH or pSBbi- AGX1$^{F383A}$-NahK, were either collected for RNA extraction (unfed) or fed with DMSO, 10µM $Ac_4$GalN6yne or 10µM $Ac_4$GalNAc. After overnight feeding, cells were collected for RNA extraction.

RNA was extracted from fed and unfed cells by using RNeasy Mini kit (Qiagen, Crawley, UK), DNase I (Thermo Fisher) and QIAshredder (Qiagen ) according to the manufacturer's instructions. mRNA capture and library preparation were performed by the Advanced Sequencing Facility at the Francis Crick Institute using the KAPA mRNA HyperPrep Kit (Roche, Basel, Switzerland). Technical triplicate libraries were sequenced on an Illumina HiSeq 4000 platform at the facility to generating on average 12 million 101 bp single-end reads per sample.

Raw reads were quality and adapter trimmed using cutadapt (version 1.5)[12] before alignment. Reads were mapped and subsequent gene-level counted using RSEM 1.3.0[13] and STAR 2.5.2[14] against the human genome GRCh38 using annotation release 86, both from Ensembl. Normalisation of raw count data and differential expression analysis was performed with the DESeq2 package (version 1.24.0)[15] within the R programming environment (version 3.6.1)[16]. The following pairwise comparison were performed: pSBbi- AGX1$^{F383A}$-NahK samples vs pSBbi-GH samples; pSBbi- AGX1$^{F383A}$-NahK-DMSO-fed samples vs GH-pSBbi-DMSO-fed samples;

pSBbi-AGX1$^{F383A}$-NahK-GalNAc-fed samples vs pSBbi-GH- GalNAc-fed samples; pSBbi-AGX1$^{F383A}$-NahK-GalN6yne-fed samples vs pSBbi-GH-GalN6yne-fed samples; and a like hood ratio test (LRT) analysis for each cell type across all feeding regimes with the contrast function, from which genes differentially expressed (adjusted p value being less than 0.05) between different conditions were determined. Gene lists were used to look for pathways, biological processes, cellular components and molecular functions enrichment using the Broad's GSEA software (version 3.0) with genesets from MSigDB (version 7.1).[17]

*SILAC-based quantitative proteomics analysis*

K-562 cells stably transfected with pSBbi-GH and pSBbi- AGX1$^{F383A}$-NahK were individually grown in *heavy* and *light* media for 6 doublings, sufficiently to achieve a labelling efficiency of >95%, before being fed with either DMSO or 10 µM $Ac_4$GalN6yne.

Light media contained RPMI (Thermo Fisher) with 10% (v/v) dialysed FBS, proline (0.1mg/mL. Thermo Fisher) and $^{12}$C/$^{14}$N light lysine/arginine K0/R0 (0.1mg/mL, Thermo Fisher) that replace normal Lys and Arg.

Heavy media contained RPMI (Thermo Fisher) with 10% (v/v) dialysed FBS, proline (0.1mg/mL) and $^{13}$C6, $^{15}$N2 heavy lysine/arginine K8/R10 (0.1mg/mL, Thermo Fisher) that replace normal Lys and Arg.

After 20 h, cells were centrifuged (500 g, 5 min) and washed twice with PBS (200 µL). After being transferred into a V-shaped 96 well plate, cells were lysed with 200 µL of ice-cold Lysis Buffer containing 50 µM PUGNAc. Cells were lysed for 20 min at 4 °C on an orbital shaker and centrifuged (1500 g, 20 min, 4 °C). Supernatant was transferred to a new plate and Pierce™ BCA Protein Assay kit (Thermo Fisher) was used to measure protein concentration.

Heavy and light lysates were mixed 1:1 (0.5 mg), normalized up to 250 µL with PBS and incubated for 2h at RT with 300 µL of Neutravidin beads slurry (Sera-Mag SpeedBeads Neutravidin-Coated Magnetic Beads, cytiva, Marlborough, USA), previously washed with PBS (2x 200 µL), to remove endogenous biotinylated proteins. The supernatant was collected, diluted to 270 µL with PBS and 30 µL of 10X CuAAC solution (3 mM CuSO4, 6 mM BTTAA, 1 mM biotin-picolyl azide (Click Chemistry Tools), 50 mM sodium ascorbate and 50 mM aminoguanidinium chloride) was added. The click reaction was left 6 h at RT under shaking. Samples were treated with 3 mL cold methanol (-20 °C, 10-fold excess) and left 24 h at -80 °C for protein precipitation.

Samples were then centrifuged (3700 g, 4 °C, 20 min) and supernatant discarded; pellets were washed twice with cold methanol (3 mL) and centrifuged between washes (3700 g, 4 °C, 20 min). Supernatant was completely removed (tubes upside-down on tissue paper, then let air dry) and samples were resuspended in 250 µL 0.1% (w/v) Rapigest (Waters) in PBS and sonicated in a water bath for 25 min. Samples were centrifuged (3700 g, 5 min), supernatants were transferred to new tubes and pellets were treated with 250 µL of 6 M urea in PBS. Samples were sonicated for 25 min and centrifuged again (3700 g, 5 min). The pellets were resuspended with 250 µL of PBS, sonicated for 25 min and centrifuged again. Rapigest, urea and PBS supernatants were then combined and incubated with 350 µL of Neutravidin Beads slurry (previously washed twice with

200 µL of PBS) for 2 h at RT. Beads were washed with 1% (w/v) Rapigest (3x 350 µL), 6 M urea in PBS (6x 350 µL), AmBic (50 mM ammonium bicarbonate, 6x 350 µL) and 40% (v/v) LCMS-grade acetonitrile (4x 100 µL). Beads were resuspended in 100 µL of AmBic containing 10 mM DTT and then incubated at 50 °C for 15 min. Beads were washed with AmBic (2x 350µL) and 100 µL of 20 mM iodoacetamide in AmBic was then added. Samples were kept for 30 min in the dark. Iodoacetamide was then quenched by adding DTT 10 mM (final concentration). The beads were washed with AmBic (3x 350µL), then resuspended in 100 µL of AmBic and 300 ng of LysC Mass Spec Grade (Promega) were added to beads followed by overnight incubation at 37 °C. The supernatant was transferred to a new tube and 200 ng of trypsin gold Mass Spec Grade (Promega) were added. The digestion was left for 8 h at 37 °C. Peptides were desalted by UltraMicroSpin™ (The Nest group Inc., Ipswich, USA) according to the manufacturer protocol and vacuum-dried by SpeedVac to remove any traces of organic solvents.

Dried peptides were resuspended in 16 µL of 0.1% (v/v) formic acid in LCMS-grade water, sonicated for 15 min in water-bath, vortexed briefly and harvested 5 min at 18,000 g. Peptide mixtures were analysed by nanoflow LC-MS/MS using an Orbitrap Fusion Lumos with ETD (Electron Transfer Dissociation Mass Spectrometry, Thermo Fisher) coupled to an UltiMate 3000 RSLCnano (Thermo Fisher). The sample (15 µL) was loaded via autosampler isocratically onto a 50 cm, 75 µm PepMap RSLC C18 column after pre-concentration onto a 2 cm, 75 µm Acclaim PepMap100 m nanoViper.

The column was held at 40 °C using a column heater in the EASY-Spray ionization source (Thermo Fisher). The samples were eluted at a constant flow rate of 0.275 µL/min using a 240 minutes gradient. Solvents were: A = 5% (v/v) DMSO, 95% (v/v) 0.1% formic acid in water; B = 5% (v/v) DMSO, 20% (v/v) 0.1% formic acid in water, 75% (v/v) 0.1% formic acid in acetonitrile. The gradient profile was as follow: 0 min 98% A, 2% B; 5 min 98% A, 2% B; 55 min 98% A, 2% B; 190 min 70% A, 30% B; 213 min 60% A, 40% B; 213 min 5% A, 85% B; 213 min 5% A, 85% B; 225 min 98% A, 2% B: 240 min 98% A, 2% B.

MS1 scans were collected with a mass range from 350-1500 m/z, 120K resolution, 4e5 ion inject target, and 50 ms maximum inject time. Dynamic exclusion was set to exclude for 45 seconds with a repeat count of 1. Charge states 2-6 with an intensity greater than 1E4 were selected for fragmentation at top speed for 3s. Selected precursors were fragmented using HCD at 30% nCE with 1.2 Da isolation window, 1e4 inject target, and 100 ms maximum inject time. MS2 scans were taken in the ion trap at a rapid scan rate.

Raw mass spectrometry files were loaded into MaxQuant software[18] for quantification and identification by using *homo sapiens* FASTA protein sequences database from UniProt (downloaded 18 June, 2020) for the database search.[19] The protein groups table were uploaded into Perseus[20] to allow for data transformation and visualization, and into RStudio for statistical analysis.

Search parameters included standard group-specific parameter type with multiplicity of 2 and maximum labelled of 3. Specific cleavage specificity of R and K, with two missed cleavages were allowed. Methionine oxidation and N-terminal acetylation were set as variable modifications with a total common max of 5. Cysteine carbamidomethylation was set as a fixed modification. Peptide hits were filtered using a 1% false-discovery rate.

*Fluorescence microscopy*

For mono-culture samples, non-transfected and pSBbi-AGX1$^{F383A}$-NahK stably transfected 4T1(GFP-expressing) and fibroblast MLg cells were seeded into a µ-Plate 24 Well Black (Thistle Scientific Ltd, Glasgow, UK) at a density of 30,000 cells in 350 µL growth medium without hygromycin. Cells were treated with either DMSO, 50 µM $Ac_4$GalN6yne or 25 µM $Ac_4$ManAlk.

For co-culture samples, non-transfected and pSBbi-AGX1$^{F383A}$-NahK stably transfected MLg cells were seeded into a µ-Plate 24 Well Black (Thistle Scientific Ltd) at a density of 30,000 cells in 350 µL growth medium without hygromycin. After 4 h, 3000 4T1(GFP) cells were plated on the top of the MLg cells (1:10 ratio). The co-culture samples were grown 72 h before feeding with either DMSO, 50 µM $Ac_4$GalN6yne or 50 µM $Ac_4$ManAlk.

Cells were incubated for 16 h. Medium was aspirated, and cells were washed with ice-cold 2% (v/v) FBS in PBS (2x 200 µL). Cells were then treated with 200 µL of a freshly prepared solution containing 300 µM BTTAA, 50 µM $CuSO_4$, 5 mM sodium ascorbate, 5 mM aminoguanidinium chloride and 200 µM biotin-picolyl azide. The reaction was carried out for 3 min at room temperature, the supernatant was aspirated and cells were washed with ice-cold PBS (4x 200 µL). Cells were incubated for 20 min at RT with 20 µg/mL Streptavidin-AlexaFluor647 (BioLegend UK Ltd, Kentish Town, UK) in 1% (w/v) BSA in PBS solution in the dark. After washing with ice-cold PBS (4x 200 µL), cells were fixed 20 min with cold 4% (v/v) formaldehyde (Thermo Fisher) in 100 mM sodium phosphate buffer pH 7.4, at room temperature in the dark. The reaction was quenched by 5 min incubation with 50 mM ammonium chloride and the cells were washed with PBS (3x 200 µL). Cells were permeabilized with 0.1% (v/v) Triton X-100 in PBS for 10 min at 4 °C and washed with PBS (3x 200 µL). Cells were blocked in a solution of 10% (v/v) normal donkey serum (ab7475, abcam), 1% (w/v) BSA and 0.1% (v/v) Tween-20 in PBS for 1 h at room temperature. GFP expression was detected by incubation with goat anti-GFP (1:300, ab5450, abcam) in a solution containing 5% (v/v) donkey serum in 1% (w/v) BSA in PBS. Cells were washed with PBS (3x 200 µL) and incubated for 30 min at RT with Alexafluor488 anti-goat secondary antibody (1:500, ab150129, abcam) in a solution containing 1% (v/v) normal donkey serum in 1% (w/v) BSA in PBS. Cells were then incubated with Alexafluor568 Phalloidin (Invitrogen A12380, 5 µL of 40X methanol stock solution in each 200 µL of PBS) in 1% (w/v) BSA and 0.1% (v/v) Tween-20 followed by PBS washing (3x 200 µL) and DAPI incubation (1:1000, Vector Laboratories Ltd, Peterborough, UK) in 1% (w/v) BSA and 0.1% (v/v) Tween-20 in PBS (Thermo Fisher) for 30 min at room temperature. After washing with PBS (3x 200 µL), circle CoverSlips with 15 mm diameter (Thermo Fisher) were mounted onto each well by using 20 µL of ProLong Gold antifade reagent (Invitrogen).

The confocal acquisition was made on a Zeiss LSM710 Invert microscope. The images were acquired using a Plan Apochromat 40x/1.3 Oil objective. Mono-culture samples were imaged with an acquisition zoom of 1.3, so the corresponding resulting pixel size was 0.159 µm. For the co-culture samples, three-dimensional images were acquired with an acquisition zoom of 0.6, so the corresponding resulting pixel size was 0.219 µm (x, y) and 0.7 µm (z).
A sequential scan to spectrally separate the fluorescence of DAPI, Alexafluor 647, Alexafluor488 and Alexafluor568 was used. In addition, the transmitted light channel was activated to visualize the cell morphology. Images were visualized and processed with Fiji and Zen (Zeiss, Oberkochen, Germany) software.[21]

*Proteomics and glycoproteomics analysis in co-culture samples of secretome and cell lysate*

Murine 4T1(GFP-expressing) and human MCF7 stably transfected with pSBbi-AGX1$^{F383A}$-NahK-T2$^{BH}$ and pSBbi-Hyg plasmids were individually plated or co-cultured (1:1 ratio) overnight in DMEM with 10% (v/v) FBS, penicillin (100 U/mL), streptomycin (100 µg/mL) and hygromycin B (100 µg/mL). For each mono- and co-culture sample, two 15 cm petri dishes containing a total of $6 \times 10^6$ cells were prepared.
After overnight growing, the media was discarded and replaced with fresh media containing either DMSO or 10 µM $Ac_4GalN6yne$. FBS-free media was used for feeding samples addressed to secretome glycoproteins labelling.
**For secretome glycoprotein enrichment**, media was collected and centrifuge (500 g, 5 min) to remove any cell debris. The secretome was then concentrated up to 300 µL and media replaced with PBS by using an Amicon Ultra-15 Centrifugal Filter Unit (3 kDa MWCO, Merck). Pierce™ BCA Protein Assay kit (Thermo Fisher) was used to measure protein concentration. 300 µg of each sample was used for the next step.
**For the cell lysate glycoproteins enrichment**, media was discarded and cells were washed with PBS (5 mL). Cells were detached from the petri dish by 10 min incubation with 8 mM EDTA in PBS and then washed twice with PBS. Cells were lysed with 200 µL of ice-cold Lysis Buffer containing 50 µM PUGNAc. Cells were lysed for 20 min at 4 °C on an orbital shaker and centrifuged (1500 g, 20 min, 4 °C). Supernatant was transferred to a new plate and Pierce™ BCA Protein Assay kit (Thermo Fisher) was used to measure protein concentration.
For each lysate (secreotome or lysate), 300 µg of protein was brought to 250 µL with PBS and incubated for 2 h at RT with 300 µL of Neutravidin beads slurry (Sera-Mag SpeedBeads Neutravidin-Coated Magnetic Beads, cytiva, Marlborough, USA), previously washed with PBS (2x 200 µL), to remove endogenous biotinylated proteins.
Both secretome and lysate samples were diluted to 270 µL with PBS and 30 µL of 10X CuAAC solution (3 mM CuSO4, 6 mM BTTAA, 1 mM DADPS-biotin-azide (Click Chemistry Tools), 50 mM sodium ascorbate and 50 mM aminoguanidinium chloride) were added. The click reaction was left 6 h at RT under shaking. Samples were treated with 3 mL cold methanol (-20 °C, 10-fold excess) and left 24 h at -80 °C for protein precipitation.

Glycoproteins were enriched on Neutravidin Beads by following the same procedure described in the *SILAC-based quantitative proteomics analysis* section. Peptides were eluted as described in the above mentioned section.

Beads with bound glycopeptides following on-bead digest were incubated with 150 µL of 1% (v/v) formic acid in LCMS-grade water 30 min at RT on a rotator. The supernatant was collected and the acid-cleavage treatment was repeated a second time. Beads were washed with LCMS-grade acetonitrile. The wash and the acidic supernatants were combined together and 200 ng of trypsin were added. The digestion was left 8 h at 37 °C. Glycopeptides were dried by SpeedVac and then resuspended in AmBic containing 2% (v/v) LCMS-grade acetonitrile and 0.1% (v/v) formic acid.

Peptides and glycopeptides were desalted by UltraMicroSpin$^{TM}$ (The Nest group Inc., Ipswich, USA) according to the manufacturer protocol and vacuum-dried by SpeedVac to remove any traces of organic solvents.
Dried peptides and glycopeptides were resuspended in 16 µL of 0.1% (v/v) formic acid in LCMS-grade water, sonicated for 15 min, vortexed briefly and centrifuged for 5 min at 18,000 g.
Sample mixtures were analysed by nanoflow LC-MS/MS using an Orbitrap Eclipse with ETD (Thermo Fisher) coupled to an UltiMate 3000 RSLCnano (Thermo Fisher).
The sample (15 µL for glycopeptide fractions and 5µL out of 16 µL for peptide fractions) was loaded via autosampler isocratically onto a 50 cm, 75 µm PepMap RSLC C18 column (ES903) after pre-concentation onto a 2 cm, 75 µm Acclaim PepMap100 m nanoViper.
The column was held at 40 °C using a column heater in the EASY-Spray ionization source (Thermo Fisher). The samples were eluted at a constant flow rate of 0.275 µL/min using a 120 and 140 minutes gradient for peptides and glycopeptides, respectively. Solvents were: A = 5% (v/v) DMSO, 95% (v/v) 0.1% formic acid in water; B = 5% (v/v) DMSO, 20% (v/v) 0.1% formic acid in water, 75% (v/v) 0.1% formic acid in acetonitrile.
**For the peptides,** the gradient profile was as follows: 0 min 98% A, 2% B; 5 min 98% A, 2% B; 5.5 min 92% A, 8% B; 93 min 60% A, 40% B; 94 min 5% A, 95% B; 104 min 5% A, 95% B; 105 min 98% A, 2% B; 120 min 98% A, 2% B.
**For the glycopeptides,** the gradient profile was as follows: 0 min 98% A, 2% B; 6 min 98% A, 2% B; 114 min 60% A, 40% B; 115 min 95% A, 5% B; 119 min 5% A, 95% B; 120 min 98% A, 2% B: 140 min 98% A, 2% B.

**For unmodified peptide identification,** MS1 scans were collected with a mass range from 350-1500 m/z, 120K resolution, 4e5 ion inject target, and 50 ms maximum inject time. Dynamic exclusion was set to exclude for 20 seconds with a repeat count of 1. Charge states 2-6 with an intensity greater than 1e4 were selected for fragmentation at top speed for 3s. Selected precursors were fragmented using HCD at 30% nCE 1.2 Da isolation window, 1e4 inject target, and 100 ms maximum inject time. MS2 scans were taken in the ion trap at a rapid scan rate.

**For glycopeptide identification,** MS1 scans were collected with a mass range from 300-1500 m/z, 120K resolution, 4e5 ion inject target, and 50 ms maximum inject time. Dynamic exclusion was set to exclude for 10 seconds with a repeat count of 3. Charge states 2-6 with an intensity greater than 1e4 were selected for fragmentation at top speed for 3s. Selected precursors were fragmented using HCD at 28% nCE with 2 Da isolation window, 5e4 inject target, and 54 ms maximum inject time before collection at 30K resolution in the Orbitrap. For precursors from 300-1000 m/z, presence of 3 oxonium ions over 5% relative abundance triggered a charge calibrated ETD scan to be collected in the ion trap with a 3 Da isolation window, 1e4 inject target, and 100ms maximum injection time.

**For unmodified peptide analysis,** raw mass spectrometry files were loaded into MaxQuant software[18] for quantification and identification by using Homo sapiens (downloaded 18 June, 2020) and Mus musculus (downloaded 10 September, 2020) FASTA protein sequences database from UniProt as a reference database. For peptides, search parameters included specific cleavage specificity of R and K, with two missed cleavages allowed. Methionine oxidation and N-terminal acetylation were set as variable modifications with a total common max of 5. Carbamidomethyl cysteine was set as a fixed modification. Peptide hits were filtered using a 1% FDR. The protein groups table were uploaded into Perseus[20] to allow for data transformation, visualization and statistical analysis.

**Data evaluation of glycopeptides** was performed with Byonic™ (Protein Metrics, Cupertino, USA). For glycopeptide analysis, search parameters included semi-specific cleavage specificity at the C-terminal site of R and K, with two missed cleavages allowed. Mass tolerance was set at 10 ppm for MS1s, 20 ppm for HCD MS2s, and 0.2 Da for ETD MS2s. Carbamidomethyl cysteine was set as a fixed modification. Variable modifications included methionine oxidation (common 1), asparagine deamidation (common 1), and a custom database of O-glycans that included HexNAc, HexNAc-NeuAc, HexNAc-Hex, HexNAc-Hex-NeuAc, HexNAc2-Hex-NeuAc and HexNAc-Hex-NeuAc2 with an additional 287.1371 m/z to account for the chemical modification. A maximum of two variable modifications were allowed per peptide. All identifications with |logP| greater than 3 that contained chemically modified glycans were manually validated and localized using a combination of HCD and ETD information.

*Mouse experiments*

NOD-SCID IL2Rgnull (NSG) strain mice were obtained from the Jackson Laboratory and bred at the Francis Crick Institute Biological Resources Facility in individually vented cages under specific-pathogen-free (SPF) conditions. When performing the mammary fat pad injection of cancer cells into the mice, the mice were anaesthetized with inhaled isoflurane.

All animals in the experiments discussed here were performed under project license (P83B37B3C), approved by the UK Home Office, and in accordance with The Francis Crick Institute animal ethics committee guidelines.

*Bioorthogonal cell-specific tagging of glycoproteins in vivo.*

GFP-expressing 4T1 cells transfected with pSBbi-AGX1$^{F383A}$-NahK-T2$^{BH}$ were resuspended in 50 µL of growth-factor-reduced Matrigel (BD Biosciences) and injected into the left hind fat pad of female NSG mice, with GFP-expressing 4T1 cells transfected with empty pSBbi-Hyg injected into the right hind fat pad of the mouse as an internal control (1x10$^{6}$ cells per injection). Body weight and tumour volume were monitored every other day.

**For intratumoural compound administration:** 11 days post injection, mice harbouring tumours of roughly 200-300 mm$^{3}$ were randomly assigned to receive intratumoral injection with 50 µL of vehicle (5% (v/v) DMSO/PEG-400; n = 1) or $Ac_4$GalN6yne (6 mg/mL in 5% (v/v) DMSO/PEG-400; n = 2) once daily for three consecutive days. On the fourth day the tumours were harvested, flash-frozen and homogenized for Western blot analysis. In the case of the BOCTAG-T2 tumours, an additional piece was prepared for tumour digestion, protein expression analysis and metabolic cell surface labelling (see below for procedures).

**For intraperitoneal compound administration:** 13 days post tumour cell injection, mice were randomly divided into three groups and intraperitoneally injected with 100 µL of vehicle (5% (v/v) DMSO/PEG-400; n=1), $Ac_4$GalN6yne (40 mg/mL in 5% (v/v) DMSO/PEG-400; n = 3) or $Ac_4$ManNAlk (40 mg/mL in 5% (v/v) DMSO/PEG-400; n = 2) once daily for five consecutive days. On the sixth day, the tumours were harvested, flash-frozen and homogenized for Western blot analysis. In the case of the BOCTAG-T2 tumours, an additional piece was prepared for tumour digestion, protein expression analysis and metabolic cell surface labelling (see below for procedures).

**Tumour homogenization and Western blot analysis.** Tumour pieces were homogenized in ice-cold Lysis Buffer containing 50 µM of PUGNAc using a Precellys© homogeniser (speed = 4000 g, number of cycles = 3, cycle duration = 30 s, waiting time between 2 cycles = 30 s, temperature = 4 °C). The samples were allowed to stand for 10 min and subsequently centrifuged (10000 g, 4 °C, 15 min) to remove cell debris. The supernatant was collected and the protein concentration determined by BCA assay.
To remove endogenous biotinylated proteins, the samples were diluted in PBS to a protein content of 3.3 mg/mL. Neutravidin-coated magnetic beads (375 µL slurry = 175 µL settled resin), previously washed with PBS (3x 200 µL), were added to the samples and incubated for 2 h at RT under rotation. The supernatant was subsequently collected and the protein concentration determined by BCA assay.
The collected supernatant (30 µg protein) was then treated with a freshly prepared solution containing 600 µM BTTAA, 300 µM $CuSO_4$, 5 mM sodium ascorbate, 5 mM aminoguanidinium chloride and 100 µM biotin picolyl azide and incubated overnight at RT on an orbital shaker.
Loading buffer was subsequently added to the samples and these were run on a 4–20% Criterion™ TGX™ Precast gel for SDS-PAGE. After membrane transfer, the total protein amount was

assessed using the REVERT protein staining kit and biotinylation detected using IRDye 800CW Streptavidin according to the manufacturer's instructions.

**Tumour digestion, protein expression analysis and metabolic cell surface labelling of the BOCTAG samples.** The BOCTAG-T2 tumours were manually minced with a scalpel and scissors until they became a smooth paste with no visible clumps and then incubated with 1.5 mL of a freshly prepared digestion solution (a 1:3:3:193 mixture of DNase I, Liberase TM, Liberase TM and HBSS without $CaCl_2$ or $MgCl_2$) for 2h at 37 °C. The cell suspension was then filtered using a 100 µm cell strainer and the reaction quenched by adding an equal volume of growth medium (DMEM with 10% (v/v) FBS, penicillin (100 U/mL), streptomycin (100 µg/mL)) to the filtered cell suspension. The samples were subsequently centrifuged (300 g, 4 °C, 10 min) and the pellet resuspended in 10 mL growth medium. The cell suspension was then plated on a 10-cm dish and incubated overnight at 37 °C. After 24 h, the cells were washed once with PBS and treated with fresh growth medium. The cells were subsequently propagated in fresh growth medium with 50 µg/mL hygromycin B at 37 °C with 5% $CO_2$ for 10 days.

For protein expression analysis and metabolic cell surface labelling, the cells were seeded at a density of 500,000 cells/mL in 1.6 mL growth medium without hygromycin into 6-well plates and allowed to attach overnight. The cells were then treated with 50 µM $Ac_4$GalN6yne or 50 µM $Ac_4$ManNAlk or a corresponding volume of DMSO. After 20 h, the cells were washed once with PBS (without $CaCl_2$ or $MgCl_2$) and incubated with 4 mM EDTA in PBS (1.5 mL) for 5 min. The cells were transferred to a 1.5 mL centrifuge tube, harvested (500 g, 4 °C, 5 min) and resuspended in ice-cold Labelling Buffer (0.2 mL). The metabolic cell surface labelling and in-gel fluorescence analysis was subsequently performed as explained above.

Protein expression was assessed by Western blot using antibodies against FLAG tag (rabbit anti-FLAG antibody,1:1000, PA1-984B, invitrogen, Carlsbad, USA), HA tag (rabbit anti-HA antibody, 1:1000, ab9110, abcam), VSV-G tag (goat anti-VSV-G, 1:2000, ab3861, abcam) and GADPH (rabbit anti-GAPDH, 1:5000, ab181602, abcam).

*Chemical synthesis*

**2-deoxy-2-(5-hexynoyl)amido-α-D-galactopyranosyl phosphate disodium salt (SI-3)**

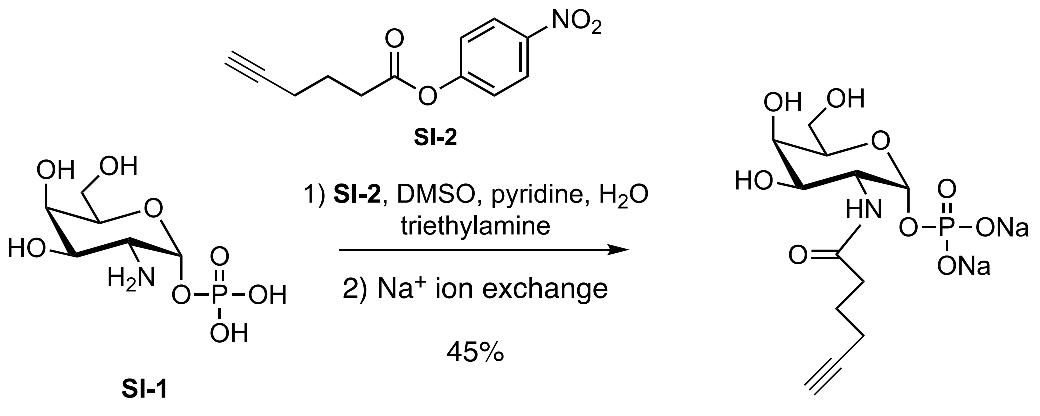


GalN6yne-1-phosphate (**SI-3**)

Amine **SI-1** (15 mg, 58 µmol) in anhydrous DMSO/pyridine (1:1.5 (v/v), 1.2 mL) was treated with 4-nitrophenyl hex-5-ynoate **SI-2** (20 mg, 87 µmol) and triethylamine (8 µL, 58 µmol). The turbid solution was left to stir overnight. The reaction was treated with 100 µL water and left to stir for another 8 h. Another 150 µL water were added as well as 1.5 equiv. reagent **SI-2** (20 mg, 87 µmol). After another 60 h, another 200 µL water were added, then another 1.5 equiv. reagent **SI-2** (20 mg, 87 µmol). The yellow solution turned clear overnight, when TLC (DCM/MeOH/water 4:1:1 with 2 drops triethylamine) indicated conversion. The solution was shock frozen on dry ice, lyophilized and purified by size exclusion chromatography (Sephadex G-25 extra fine, Sigma-Aldrich) using water/MeOH 3:2 (v/v) as a solvent. The combined fractions were concentrated and passed through AG 50W-X8 $Na^+$ form resin (Bio-Rad) and lyophilized to give alkyne **SI-3** (10 mg, 26 µmol, 45%) as a white solid. $^1$H NMR (600 MHz, $D_2O$) δ 5.40 (dd, *J* = 7.5, 3.5 Hz, 1H), 4.31 – 4.14 (m, 2H), 4.01 (s, 1H), 3.94 (dd, *J* = 10.9, 3.1 Hz, 1H), 3.84 – 3.67 (m, 2H), 2.46 (m, *J* = 14.8, 7.2 Hz, 2H), 2.39 – 2.33 (m, 2H), 2.28 (m, 4H), 2.23 (m, 1H), 1.84 (m, 1H), 1.76 (m, 1H); $^{13}$C NMR (150 MHz, $D_2O$) δ 177.1, 93.1, 71.4, 71.2, 70.2, 69.6, 69.1, 68.9, 68.4, 61.6, 50.4, 36.7, 34.9, 24.9, 24.4, 17.3; HRMS (ESI) calcd. for $C_{12}H_{20}NO_9P$ (M-$H^+$) 353.0876 found 352.0800 *m/z*.

$^{1}H$ NMR (600 MHz, $D_2O$)

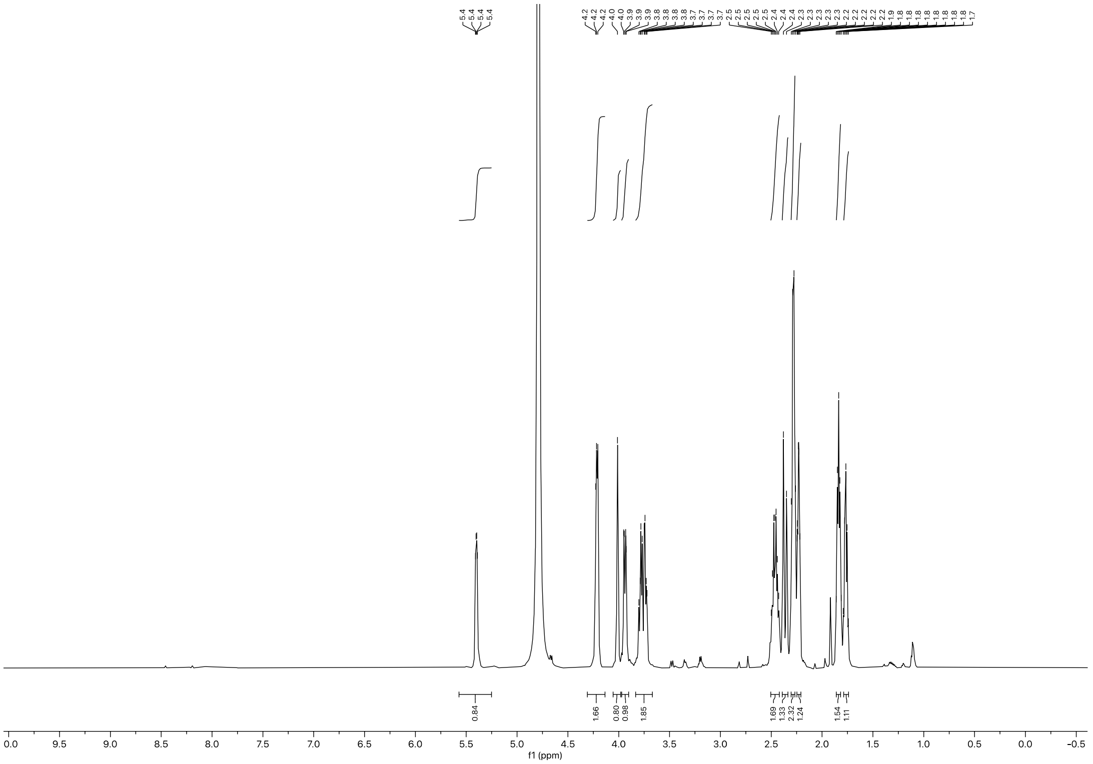


$^{13}C$ NMR (150 MHz, $D_2O$)

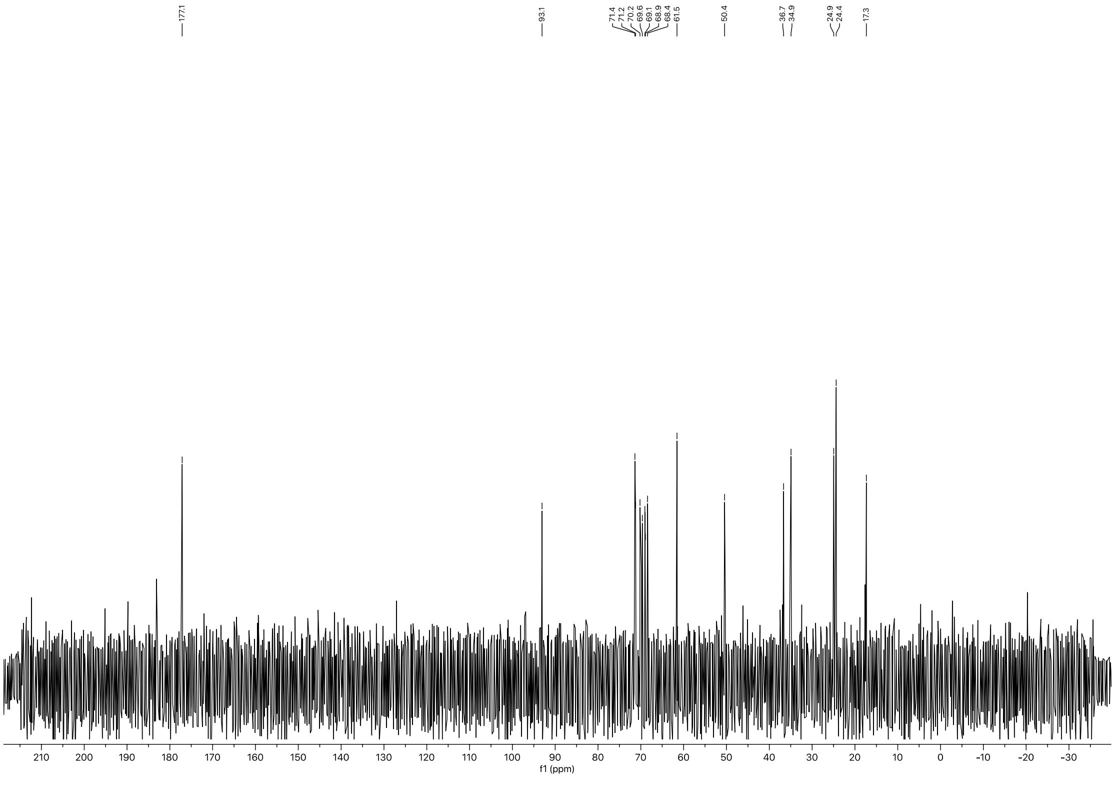

### Appendix:

Sequence of pSBbi-AGX1$^{F383A}$-NahK-T2$^{I253A/L310A}$

**GGATCCCTATACAGTTGAAGTCGGAAGTTTACATACACTTAAGTTGGAGTCATTAAAACT
CGTTTTTCAACTACTCCACAAATTTCTTGTTAACAAACAATAGTTTTGGCAAGTCAGTTAG
GACATCTACTTTGTGCATGACACAAGTCATTTTTCCAACAATTGTTTACAGACAGATTATT
TCACTTATAATTCACTGTATCACAATTCCAGTGGGTCAGAAGTTTACATACACTAAGTTCG
ACTCCTCTGCAGAATGCGGCGATGTTTCGGTAAGGGGTCCGCTATCTAGTAGGCCCAGC
TGGTTCTTTCCGCCTCAGAAGCCATAGAGCCCACCGCATCCCCAGCATGCCTGCTATTGT
CTTCCCAATCCTCCCCCTTGCTGTCCTGCCCCACCCCACCCCCCAGAATAGAATGACACC
TACTCAGACAATGCGATGCAATTTCCTCATTTTATTAGGAAAGGACAGTGGGAGTGGCAC
CTTCCAGGGTCAAGGAAGGCACGGGGGAGGGGCAAACAACAGATGGCTGGCAACTAGA
AGGCACAGTCGAGGCTGATCAGCGAGCTCTAGAGAATTGATCCCCAAGCTTGGCCTGAC
AGGCCCTACTTACCCAGGCGGTTCATTTCGATATCAGTGTACTGCTGCAGGTTGAGCGTG
AACTTCCACTGCTGCGAAAGGGCCGGGCCACACACCTCCACGCTTAGGCCCCCGCTCTT
GGCCGTGCGACTGTCCAGGCACAGGTTGCTGCCCACGTGCCTCAGCTTGGAGTTGCCCT
CGATCTGTTCCCATTTCTGTCTGCTGTCATTTTCTCGGCAGCCCTGCAGCTTTATAAGAG
AGCCCGGTGCCCGGTCCACCACAGTAAGGCACAAATCCATGTGCTTCACCGACTTCTCCT
TCGTCAAGGCCCATTCCTGGTTTCCCCCAGCATTGTGACATTCATAAACTCCAACCACAC
CATCAGCAAAGTGTCCCAAAGTGTCGAGGCAGTTAGTTCCCTGCTGCAAGGCCCCAAAA
GCTATATCCTGATGGTCTGGAACCCTTAACTCTGGATAGACATTTTCAAGGTACCATTTG
AAAGGCTTGCAGCTGAGTTTCTTCCTAAGCTCCAATCTGCTCTGAATATTTCCATAAGGA
ACGTTTCTAGCAGAAGGCACTGCTGCATAATAGAAATTTTTGTATTCATCCATCCAGACC
TCTGCTGCCCGGCGGGTGTTTCGGGCAAAGACAGTGCCACTGCCACCCGGGAACGTGTA
GGGGTGCTGCTTCCGGAACACGTGTCCCACACGGCTGCACGGGATGATCTCCAGGCTGC
CACCACACTGCCACACGCGGAACGAGATCTCTAGGTTCTCTCCTCCCCACACATCCATCA
TCATGTCGTACTTCCCCAGTTCTTCAAAATAGAACTTATCCATCACAAAGGCCCCACCAG
CAATCATGGGGGTTTTTATAGGGGCGACTGGGTTCCCCTGCCGGGACCTTCTCTGCTCAG
GCGTCATGTAATCCCACTTGAATACCAAGTTCCAATCAAAACCGCCCTTCAAGTCAGCAG
ATGCCCCCACATACTGAAAGTTGTCCATATTAATGACATCGGCGATGGGTGACACAACCC
GAGTCCTGTCCTCCGCCACCCTTTCCAGGAGGGGCTCCAGCCAGTGCTCATTACACTCGC
AGTGACTGTCCAGGAAGGTCAGGACCTTGGCTTGGGCAGCATCGGCCCCCCGAACCCGT
GAGCGCATGAGGCCTTCTCGTCGATCATTTCTAAGAACTCGCACTTTCTCAATTTTCCCC
AAGAGAGCCCCGTCCTCAGGATCATTGCTGTAGTCATCCACCAAGATGATTTCTTTTATG
AGATGGGGCGGGCTTTTCTTAAGCACGCTGACCACGGTCCTGAGTAGGGCCGACCTGGC
TTCATTGTGAAACGTGATCACCACGCTGGTGGCCGGCAGATCCACCCGCCACTGCTTCCG
CTGACACTGGTCATGCCGGGTGTCAGGGATGGCTCTGTCCATTCGAAGCTTATCACTCTC
CACCTGGTTGAACTTGTTGCGGGCGTAAGGGTCCTGCCCGGAGCGGACCATCGTCCCTC
CAACATAAGCTTCCTGGTTAAAGTCTGGCCACCGTACTTTCCCTGGAGGGAGGGTCTCCA
TGCTTTGTGCTTTCTCTTCTCCATTGCTGTGATGAAGGTCTTTCTTTTTAATGGGGTCAAT
TTCATTCCAGTCCTCCTTCCTGCCGGCGCCGCCGCCCGCGCCCCCGGCCAGCGCAGAGC
CGCCCCCCGAGTACATGTAGTAGGCGATGCCCAGCACCCACAGGAAGGCGAAGCAGAGC
AGCATCCGCGAGCGCCGCCGCATGGTGGCCTCAGAGGCCAGCTTGGGGTAGTTTTCACG
ACACCTTAAATGGAAGAAAAAAACTTTGAACCACTGTCTGAGGCTTGAGAATGAACCAAG
ATCCAAACTCAAAAAGGGCAAATTCCAAGGAGAATTACATCAAGTGCCAAGCTGGCCTAA
CTTCAGTCTCCACCCACTCAGTATGGGGAAACTCCATCGCATAAAACCCCTCCCCCCAAC
CTAAAGACGACGTACTCCAAAAGCTCGAGAACTAATCGAGGTGCCTGGACGGCGCCCGG
TACTCCGTGGAGTCACATGAAGCGACGGCTGAGGACGGAAAGGCCCTTTTCCTTTGTGT
GGGTGACTCACCCGCCCGCTCTCCCGAGCGCCGCGTCCTCCATTTGAGCTCCCTGCAG
CAGGGCCGGGAAGCGGCCATCTTTCCGCTCACGCAACTGGTGCCGACCGGGCCAGCCTT
GCCGCCCAGGGCGGGGCGATACACGGCGGCGCGAGGCCAGGCACCAGAGCAGGCCGGC
CAGCTTGAGACTACCCCCGTCCGATTCTCGGTGGCCGCGCTCGCAGGCCCCGCCTCGCC
GAACATGTGCGCTGGGACGCACGGGCCCCGTCGCCGCCCGCGGCCCCAAAAACCGAAAT**

ACCAGTGTGCAGATCTTGGCCCGCATTTACAAGACTATCTTGCCAGAAAAAAAGCGTCGC
AGCAGGTCATCAAAAATTTTAAATGGCTAGAGACTTATCGAAAGCAGCGAGACAGGCGC
GAAGGTGCCACCAGATTCGCACGCGGCGGCCCCAGCGCCCAGGCCAAGCCTCAACTCAA
GCACGAGGCGAAGGGGCTCCTTAAGCGCAAGGCCTCGAACTCTCCCACCCACTTCCAAC
CCGAAGCTCGGGATCAAGAATCACGTACTGCAGCCAGGGGCGTGGAAGTAATTCAAGGC
ACGCAAGGGCCATAACCCGTAAAGAGGCCAGGCCCGCGGGAACCACACACGGCACTTAC
CTGTGTTCTGGCGGCAAACCCGTTGCGAAAAAGAACGTTCACGGCGACTACTGCACTTAT
ATACGGTTCTCCCCCACCCTCGGGAAAAAGGCGGAGCCAGTACACGACATCACTTTCCCA
GTTTACCCCGCGCCACCTTCTCTAGGCACCGGTTCAATTGCCGACCCCTCCCCCCAACTT
CTCGGGGACTGTGGGCGATGTGCGCTCTGCCCACTGACGGGCACCGGAGCCTCTAGGCG
AGACCCTGTCTCACAAAATAAAGTAAGCCCGGACTGAGTGCGGAAAGGCGGGCCTGGCG
GGTCTGGTCTCCCCATGCGGGCCACCAGAGGCCCTGCAGCCTTCAGTCGCTTGAAGGGG
TAATGGCGCTTCCACTCACAAACATGGCGGACAGAGCGTGTGAACGAGATGAACAGCCC
CTCAAAAATATGGCCGCCGAGGCTGGACGGCCGTGCCCCAGCAGCACCGCCTCCGCGCC
CCACGTGATCTCTCGCCGGGCACAGCGCTGACCGCGGAGGTCCAACCGGAAGAATGTCC
GGATTGGACATTCGGAAGAGGGCCCGCCTTCCCTGGGGAATCTCTGCGCACGCGCAGAA
CGCTTCGACCAATGAAAACACAGGAAGCCGTCCGCGCAACCGCGTTGCGTCACTTCTGC
CGCCCCTGTTTCAAGGTATATAGCCGTAGACGGAACTTCGCCTTTCTCTCGGCCTTAGCG
CCATTTTTTTGGGTGAGTGTTTTTTGGTTCCTGCGTTGGGATTCCGTGTACAATCCATAG
ACATCTGACCTCGGCACTTAGCATCATCACAGCAAACTAACTGTAGCCTTTCTCTCTTTCC
CTGTAGAAACCTCTGCACCTGAGGCCACCATGAACATTAATGACCTCAAACTCACGTTGT
CCAAAGCTGGGCAAGAGCACCTACTACGTTTCTGGAATGAGCTTGAAGAAGCCCAACAG
GTAGAACTTTATGCAGAGCTCCAGGCCATGAACTTTGAGGAGCTGAACTTCTTTTTCCAA
AAGGCCATTGAAGGTTTTAACCAGTCTTCTCACCAAAAGAATGTGGATGCACGAATGGAA
CCTGTGCCTCGAGAGGTATTAGGCAGTGCTACAAGGGATCAAGATCAGCTCCAGGCCTG
GGAAAGTGAAGGACTTTTCCAGATTTCTCAGAATAAAGTAGCAGTTCTTCTTCTAGCTGG
TGGGCAGGGGACAAGACTCGGCGTTGCATATCCTAAGGGGATGTATGATGTTGGTTTGC
CATCCCGTAAGACACTTTTTCAGATTCAAGCAGAGCGTATCCTGAAGCTACAGCAGGTTG
CTGAAAAATATTATGGCAACAAATGCATTATTCCATGGTATATAATGACCAGTGGCAGAA
CAATGGAATCTACAAAGGAGTTCTTCACCAAGCACAAGTACTTTGGTTTAAAAAAAGAGA
ATGTAATCTTTTTTCAGCAAGGAATGCTCCCCGCCATGAGTTTTGATGGAAAATTATTTT
GGAAGAGAAGAACAAAGTTTCTATGGCTCCAGATGGGAATGGTGGTCTTTATCGGGCAC
TTGCAGCCCAGAATATTGTGGAGGATATGGAGCAAAGAGGCATTTGGAGCATTCATGTCT
ATTGTGTTGACAACATATTAGTAAAAGTGGCAGACCCACGGTTCATTGGATTTTGCATTC
AGAAAGGAGCAGACTGTGGAGCAAAGGTGGTAGAGAAAACGAACCCTACAGAACCAGTT
GGAGTGGTTTGCCGAGTGGATGGAGTTTACCAGGTGGTAGAATATAGTGAGATTTCCCT
GGCAACAGCTCAAAAACGAAGCTCAGACGGACGACTGCTGTTCAATGCGGGGAACATTG
CCAACCATTTCTTCACTGTACCATTTCTGAGAGATGTTGTCAATGTTTATGAACCTCAGTT
GCAGCACCATGTGGCTCAAAAGAAGATTCCTTATGTGGATACCCAAGGACAGTTAATTAA
GCCAGACAAACCCAATGGAATAAAGATGGAAAAATTTGTCGCTGACATCTTCCAGTTTGC
AAAGAAGTTTGTGGTATATGAAGTATTGCGAGAAGATGAGTTTTCCCCACTAAAGAATGC
TGATAGTCAGAATGGGAAAGACAACCCTACTACTGCAAGGCATGCTTTGATGTCCCTTCA
TCATTGCTGGGTCCTCAATGCAGGGGGCCATTTCATAGATGAAAATGGCTCTCGCCTTCC
AGCAATTCCCCGCTTGAAGGATGCCAATGATGTACCAATCCAATGTGAAATCTCTCCTCT
TATCTCCTATGCTGGAGAAGGATTAGAAAGTTATGTGGCAGATAAAGAATTCCATGCACC
TCTAATCATCGATGAGAATGGAGTTCATGAGCTGGTGAAAAATGGTATTGACTACAAAGA
CGATGACGACAAGGGCAGTGGAGCTACTAACTTCAGCCTGCTGAAGCAGGCTGGTGACG
TCGAGGAGAATCCTGGCCCCATGACCGAGAGCAACGAGGACCTGTTCGGCATCGCCAGC
CACTTCGCCCTGGAGGGCGCCGTGACCGGCATCGAGCCCTACGGCGACGGCCACATCAA
CACCACCTACCTGGTGACCACCGACGGCCCCCGCTACATCCTGCAGCAGATGAACACCA
GCATCTTCCCCGACACCGTGAACCTGATGCGCAACGTGGAGCTGGTGACCAGCACCCTG
AAGGCCCAGGGCAAGGAGACCCTGGACATCGTGCCCACCACCAGCGGCGCCACCTGGGC
CGAGATCGACGGCGGCGCCTGGCGCGTGTACAAGTTCATCGAGCACACCGTGAGCTACA

**ACCTGGTGCCCAACCCCGACGTGTTCCGCGAGGCCGGCAGCGCCTTCGGCGACTTCCAG**
**AACTTCCTGAGCGAGTTCGACGCCAGCCAGCTGACCGAGACCATCGCCCACTTCCACGA**
**CACCCCCCACCGCTTCGAGGACTTCAAGGCCGCCCTGGCGGCCGACAAGCTGGGCCGCG**
**CCGCCGCCTGCCAGCCCGAGATCGACTTCTACCTGAGCCACGCCGACCAGTACGCCGTG**
**GTGATGGACGGCCTGCGCGACGGCAGCATCCCCCTGCGCGTGACCCACAACGACACCAA**
**GCTGAACAACATCCTGATGGACGCCACCACCGGCAAGGCCCGCGCCATCATCGACCTGG**
**ACACCATCATGCCCGGCAGCATGCTGTTCGACTTCGGCGACAGCATCCGCTTCGGCGCC**
**AGCACCGCCCTGGAGGACGAGAAGGACCTGAGCAAGGTGCACTTCAGCACCGAGCTGTT**
**CCGCGCCTACACCGAGGGCTTCGTGGGCGAGCTGCGCGGCAGCATCACCGCCCGCGAG**
**GCCGAGCTGCTGCCCTTCAGCGGCAACCTGCTGACCATGGAGTGCGGCATGCGCTTCCT**
**GGCCGACTACCTGGAGGGCGACATCTACTTCGCCACCAAGTACCCCGAGCACAACCTGG**
**TGCGCACCCGCACCCAGATCAAGCTGGTGCAGGAGATGGAGCAGAAGGCCAGCGAGACC**
**CGCGCCATCGTGGCCGACATCATGGAGGCCGCCCGCTACCCCTACGACGTGCCCGACTA**
**CGCCTCCGGCGCTACTAACTTCAGCCTGCTGAAGCAGGCTGGTGACGTCGAGGAGAATC**
**CTGGTCCCATGAAAAAGCCTGAACTCACCGCGACGTCTGTCGAGAAGTTTCTGATCGAAA**
**AGTTCGACAGCGTCTCCGACCTGATGCAGCTCTCGGAGGGCGAAGAATCTCGTGCTTTC**
**AGCTTCGATGTAGGAGGGCGTGGATATGTCCTGCGGGTAAATAGCTGCGCCGATGGTTT**
**CTACAAAGATCGTTATGTTTATCGGCACTTTGCATCGGCCGCGCTCCCGATTCCGGAAGT**
**GCTTGACATTGGGGAATTCAGCGAGAGCCTGACCTATTGCATCTCCCGCCGTGCACAGG**
**GTGTCACGTTGCAAGACCTGCCTGAAACCGAACTGCCCGCTGTTCTGCAGCCGGTCGCG**
**GAGGCCATGGATGCGATCGCTGCGGCCGATCTTAGCCAGACGAGCGGGTTCGGCCCATT**
**CGGACCGCAAGGAATCGGTCAATACACTACATGGCGTGATTTCATATGCGCGATTGCTGA**
**TCCCCATGTGTATCACTGGCAAACTGTGATGGACGACACCGTCAGTGCGTCCGTCGCGC**
**AGGCTCTCGATGAGCTGATGCTTTGGGCCGAGGACTGCCCCGAAGTCCGGCACCTCGTG**
**CACGCGGATTTCGGCTCCAACAATGTCCTGACGGACAATGGCCGCATAACAGCGGTCAT**
**TGACTGGAGCGAGGCGATGTTCGGGGATTCCCAATACGAGGTCGCCAACATCTTCTTCT**
**GGAGGCCGTGGTTGGCTTGTATGGAGCAGCAGACGCGCTACTTCGAGCGGAGGCATCCG**
**GAGCTTGCAGGATCGCCGCGGCTCCGGGCGTATATGCTCCGCATTGGTCTTGACCAACT**
**CTATCAGAGCTTGGTTGACGGCAATTTCGATGATGCAGCTTGGGCGCAGGGTCGATGCG**
**ACGCAATCGTCCGATCCGGAGCCGGGACTGTCGGGCGTACACAAATCGCCCGCAGAAGC**
**GCGGCCGTCTGGACCGATGGCTGTGTAGAAGTACTCGCCGATAGTGGAAACCGACGCCC**
**CAGCACTCGTCCGAGGGCAAAGGAATAGTTCGAAGGCCTGTCGTGAATTCACTCCTCAG**
**GTGCAGGCTGCCTATCAGAAGGTGGTGGCTGGTGTGGCCAATGCCCTGGCTCACAAATA**
**CCACTGAGATCTTTTTCCCTCTGCCAAAAATTATGGGGACATCATGAAGCCCCTTGAGCA**
**TCTGACTTCTGGCTAATAAAGGAAATTTATTTTCATTGCAATAGTGTGTTGGAATTTTTTG**
**TGTCTCTCACTCGGAAGGACATATGGGAGGGCAAATCATTTAAAACATCAGAATGAGTAT**
**TTGGTTTAGAGTTTGGCAACATATGCCATATGCTGGCTGCCATGAACTAGCTACTCGGGA**
**CCCCTTACCGAAACATCGCCGCATTCTGCAGAGGAGTCGAGTGTATGTAAACTTCTGACC**
**CACTGGGAATGTGATGAAAGAAATAAAAGCTGAAATGAATCATTCTCTCTACTATTATTC**
**TGATATTTCACATTCTTAAAATAAAGTGGTGATCCTAACTGACCTAAGACAGGGAATTTTT**
**ACTAGGATTAAATGTCAGGAATTGTGAAAAAGTGAGTTTAAATGTATTTGGCTAAGGTGT**
**ATGTAAACTTCCGACTTCAACTGTATAGGGATCCGCTTCCTCGCTCACTGACTCGCTGCG**
**CTCGGTCGTTCGGCTGCGGCGAGCGGTATCAGCTCACTCAAAGGCGGTAATACGGTTAT**
**CCACAGAATCAGGGGATAACGCAGGAAAGAACATGTGAGCAAAAGGCCAGCAAAAGGCC**
**AGGAACCGTAAAAAGGCCGCGTTGCTGGCGTTTTTCCATAGGCTCCGCCCCCCTGACGA**
**GCATCACAAAAATCGACGCTCAAGTCAGAGGTGGCGAAACCCGACAGGACTATAAAGAT**
**ACCAGGCGTTTCCCCCTGGAAGCTCCCTCGTGCGCTCTCCTGTTCCGACCCTGCCGCTTA**
**CCGGATACCTGTCCGCCTTTCTCCCTTCGGGAAGCGTGGCGCTTTCTCATAGCTCACGCT**
**GTAGGTATCTCAGTTCGGTGTAGGTCGTTCGCTCCAAGCTGGGCTGTGTGCACGAACCC**
**CCCGTTCAGCCCGACCGCTGCGCCTTATCCGGTAACTATCGTCTTGAGTCCAACCCGGTA**
**AGACACGACTTATCGCCACTGGCAGCAGCCACTGGTAACAGGATTAGCAGAGCGAGGTA**
**TGTAGGCGGTGCTACAGAGTTCTTGAAGTGGTGGCCTAACTACGGCTACACTAGAAGAA**
**CAGTATTTGGTATCTGCGCTCTGCTGAAGCCAGTTACCTTCGGAAAAAGAGTTGGTAGCT**

| |
|---|
| **CTTGATCCGGCAAACAAACCACCGCTGGTAGCGGTGGTTTTTTTGTTTGCAAGCAGCAGA<br>TTACGCGCAGAAAAAAAGGATCTCAAGAAGATCCTTTGATCTTTTCTACGGGGTCTGACG<br>CTCAGTGGAACGAAAACTCACGTTAAGGGATTTTGGTCATGAGATTATCAAAAAGGATCT<br>TCACCTAGATCCTTTTAAATTAAAAATGAAGTTTTAAATCAATCTAAAGTATATATGAGTA<br>AACTTGGTCTGACAGTTACCAATGCTTAATCAGTGAGGCACCTATCTCAGCGATCTGTCT<br>ATTTCGTTCATCCATAGTTGCCTGACTCCCCGTCGTGTAGATAACTACGATACGGGAGGG<br>CTTACCATCTGGCCCCAGTGCTGCAATGATACCGCGAGACCCACGCTCACCGGCTCCAG<br>ATTTATCAGCAATAAACCAGCCAGCCGGAAGGGCCGAGCGCAGAAGTGGTCCTGCAACT<br>TTATCCGCCTCCATCCAGTCTATTAATTGTTGCCGGGAAGCTAGAGTAAGTAGTTCGCCA<br>GTTAATAGTTTGCGCAACGTTGTTGCCATTGCTACAGGCATCGTGGTGTCACGCTCGTCG<br>TTTGGTATGGCTTCATTCAGCTCCGGTTCCCAACGATCAAGGCGAGTTACATGATCCCCC<br>ATGTTGTGCAAAAAAGCGGTTAGCTCCTTCGGTCCTCCGATCGTTGTCAGAAGTAAGTTG<br>GCCGCAGTGTTATCACTCATGGTTATGGCAGCACTGCATAATTCTCTTACTGTCATGCCA<br>TCCGTAAGATGCTTTTCTGTGACTGGTGAGTACTCAACCAAGTCATTCTGAGAATAGTGT<br>ATGCGGCGACCGAGTTGCTCTTGCCCGGCGTCAATACGGGATAATACCGCGCCACATAG<br>CAGAACTTTAAAAGTGCTCATCATTGGAAAACGTTCTTCGGGGCGAAAACTCTCAAGGAT<br>CTTACCGCTGTTGAGATCCAGTTCGATGTAACCCACTCGTGCACCCAACTGATCTTCAGC<br>ATCTTTTACTTTCACCAGCGTTTCTGGGTGAGCAAAAACAGGAAGGCAAAATGCCGCAAA<br>AAAGGGAATAAGGGCGACACGGAAATGTTGAATACTCATACTCTTCCTTTTTCAATATTA<br>TTGAAGCATTTATCAGGGTTATTGTCTCATGAGCGGATACATATTTGAATGTATTTAGAA<br>AAATAAACAAATAGGGGTTCCGCGCACATTTCCCCGAAAAGTGCCACCTGATGCGGTGT<br>GAAATACCGCACAGATGCGTAAGGAGAAAATACCGCATCAGGAAATTGTAAGCGTTAAT<br>ATTTTGTTAAAATT** |
| **AGX1$^{F383A}$-FLAG**<br>***B. longum* NahK-HA**<br>**BH-GalNAc-T-2-VSV-G (reverse)** |
